# Evolution of hybrid inviability associated with chromosome fusions

**DOI:** 10.1101/2023.11.30.569355

**Authors:** Jesper Boman, Karin Näsvall, Roger Vila, Christer Wiklund, Niclas Backström

**Affiliations:** Evolutionary Biology Program, Department of Ecology and Genetics (IEG), Uppsala University, Norbyvägen 18D, SE-752 36 Uppsala, Sweden; Department of Zoology, University of Cambridge, Downing Street Cambridge CB2 3EJ, Cambridge, UK; Institut de Biologia Evolutiva (CSIC-Univ. Pompeu Fabra), Passeig Martim de la Barceloneta 37-49, 08003 Barcelona, Spain; Department of Zoology: Division of Ecology, Stockholm University, Stockholm, Sweden

**Keywords:** Speciation, Hybrid inviability, Hybrid incompatibilities, Chromosomal rearrangements, Population genomics

## Abstract

Chromosomal rearrangements, such as inversions, have received considerable attention in the speciation literature due to their hampering effects on recombination. However, less is known about how other rearrangements, such as chromosome fissions and fusions, can affect the evolution of reproductive isolation. Here, we used crosses between populations of the wood white butterfly (*Leptidea sinapis*) with different karyotypes to identify genomic regions associated with hybrid inviability. By contrasting allele frequencies between F_2_ hybrids that survived until the adult stage with individuals of the same cohort that succumbed to hybrid incompatibilities, we show that candidate loci for hybrid inviability mainly are situated in fast-evolving regions with reduced recombination rates, especially in regions where chromosome fusions have occurred. Our results show that the extensive variation in chromosome numbers observed across the tree of life can be involved in speciation by being hotspots for the early evolution of postzygotic reproductive isolation.

## Introduction

Understanding the genetic underpinnings of speciation lies at the heart of evolutionary biology (*1*). Since most novel species form as a consequence of reduced gene flow between incipient lineages within species (*1*, *2*), a crucial aspect of the speciation process is how barriers to gene flow are established. One such barrier is hybrid inviability, the reduced survival of hybrid offspring. Despite being at the core of speciation research for more than a century (*3*), most of our knowledge about the genetic basis of hybrid inviability comes from *Drosophila* (*4*). This is mainly a consequence of the difficulties of characterizing the genetic basis of inviable hybrids, leading to a disproportionate progress being made in model organisms that easily can be reared under controlled conditions (*4*). Crossing efforts in *Drosophila* and other organisms have shown that hybrid inviability conforms to the Bateson-Dobzhansky-Muller (BDMI) model, i.e. that alleles at two or more interacting genes are required for incompatibilities to manifest in hybrids (*4–6*). However, genic interaction is not the only mechanism by which hybrid incompatibilities can evolve.

In addition to the classical genic BDMIs, chromosomal rearrangements such as polyploidizations, gene duplications and inversions may form the genetic basis of hybrid incompatibilities. Polyploid hybrids for example, which are comparatively common in plants, are often fertile, but can be reproductively isolated from parental lineages (*7*). Chromosomal rearrangements resulting in underdominant karyotypes (hybrid underdominance model) have also been implicated in hybrid incompatibility (e.g. *8*–*10*), but this model has been criticized due to the limited parameter range under which it can evolve (*11–13*). Subpopulations evolving underdominant rearrangements need to be small and gene flow from neighboring larger populations needs to be restricted. Despite these harsh conditions, underdominant rearrangements have been documented in several animal systems (*9*, *14*).

Chromosomal rearrangements are believed to confer their fitness disadvantage by causing hybrid sterility but not hybrid inviability (*9*, *15*). However, non-disjunction in either mitosis or F_1_ hybrid meiosis may cause aneuploidies that lead to embryonic inviability (*16*). This would constitute a direct effect of chromosomal rearrangements on hybrid inviability. Chromosomal rearrangements may also contribute indirectly to speciation as a consequence of effects on the recombination rate (*17–19*). Recombination and selection are the two main processes that determine the mixing of parental haplotypes upon secondary contact (*2*, *20–22*). In non-recombining regions for example, haplotypes will segregate independently, allowing for divergence and evolution of reproductive isolation.

Based on whether chromosomal rearrangements are predicted to reduce recombination in both heterokaryotypes and homokaryotypes or not, they can be divided into two different categories: i) Rearrangements that reduce recombination only in heterokaryotypes may promote divergent evolution of genes located within the rearranged region, which can lead to reproductive isolation in the long term (*19*, *23–25*); ii) Rearrangements that reduce the recombination rate in both hetero- and homokaryotypes will result in increased selection on linked sites, in essence reducing the effective population size (*N_e_*) in the rearranged region. This leads to faster lineage sorting (*26*) and, consequently, shorter expected fixation times of segregating alleles (*27*). Regions with reduced recombination are also expected to accumulate less introgressed DNA, since introgressed regions containing deleterious alleles will be more effectively purged from the acceptor population (*20*, *22*, *28*). Thus, regions with low recombination rates have a higher probability to include loci associated with reproductive isolation. While previous theoretical and empirical work predominantly has focused on rearrangements that cause recombination suppression in heterokaryotypes, such as inversions (*1*, *17*, *18*, *25*, *29*, *30*), comparatively little is known about the consequences of chromosomal rearrangements that also reduce homokaryotype recombination, for example chromosome fusions (*10*, *21*, *31–33*).

Here, we investigate the genomic basis of hybrid inviability among populations of the wood white butterfly (*Leptidea sinapis*) with distinct karyotypes, using sequencing of large sets of pooled individuals (PoolSeq). *Leptidea sinapis* is an excellent model system to study the effects of chromosomal rearrangements on the evolution of hybrid inviability because it has the most extreme intraspecific chromosome number variation among all diploid eukaryotes (*34*). Cytogenetically confirmed chromosome numbers range from 2n = 57, 58 in Sweden (SWE) and 2n = 56-64 in Kazakhstan to 2n = 106, 108 in Catalonia (CAT; *34*, *35*). A pronounced cline in chromosome number stretches from Fennoscandia in the north and Kazakhstan in the east to the Iberian Peninsula in the south-west (*34*). A recent comparative revealed that the difference in karyotype structure between the SWE and CAT populations is a consequence of numerous fusions and fissions (*36*). While *L. sinapis* has extreme intraspecific karyotype variation, several other groups of butterflies show extensive interspecific chromosome number variation. For example between species in the *Leptidea* (*37*), *Polyommatus* (*38*) and *Erebia* (*39*) genera, and the tribe Ithomiini (*40*). Chromosome number variation in some of these groups are associated with increased diversification rates (*41*), indicating that rearrangements may have been involved in the establishment of reproductive barriers.

The wood white has previously been subject to studies on reproductive isolation since it is morphologically cryptic, but differs in genital morphology and chromosome number from the congenerics *L. reali* and *L. juvernica* (*37*, *42*). In addition, *L. sinapis* from SWE and CAT have been crossed to investigate reproductive isolation between these populations in general, and the role of the fissions and fusions in particular (*35*). In these crosses, no evidence for assortative mating was found and, despite the chromosome number difference between SWE and CAT, most hybrids were fertile (*35*). It has been hypothesized that fertility in F_1_ hybrids is rescued by a combination of inverted meiosis and holocentricity (*35*). Nevertheless, some meiotic pairing problems were observed in hybrids, indicating that the underdominance model cannot be rejected. In addition, hybrid breakdown occurred in the F_2_-F_4_ generations, with a viability of 42% compared to pure lines (*35*). This begs the question whether the extensive chromosome fusions and fissions among CAT and SWE *L. sinapis* are involved in hybrid inviability? Here, we (i) map the genomic underpinnings of hybrid inviability in *L. sinapis* using allele frequency differences between surviving F_2_ adults and F_2_ offspring that died during development, (ii) investigate the associations between recombination, chromosomal fissions and fusions and hybrid inviability, and (iii) infer the demographic history of the SWE and CAT *L. sinapis* populations and explore the evolution of hybrid inviability using population genomic methods.

## Results

### Equal survival of males and females

We crossed CAT (2n = 106-108) and SWE (2n =57, 58) chromosomal races of *L. sinapis*. Only males successfully eclosed after diapause in the ♀SWE x ♂CAT (n = 2) crosses, while both males and females eclosed in the ♀CAT x ♂SWE (n = 5) crosses. We further crossed eight F_1_ females with five F_1_ males. F_1_ females laid 3-126 eggs, producing 615 F_2_ offspring in total (Figure S1) The first =< 10 offspring of each female were collected to form a random pool of eggs, following Lima and Willett (*43*). We performed a hybrid survival experiment by monitoring the development of the remaining F_2_ offspring and observed an overall survival of 30% for both males and females (Figure 1B). Most F_2_ offspring died prior to hatching from the egg and the proportion of offspring surviving until the imago stage varied widely among families (Figure 1B-C). Since survival could be due to both genetic and environmental effects, we performed quantitative genetic analyses to estimate the genetic component of this trait. We observed a 38% narrow-sense heritability for survival (Tables S1-2 and Figure S2). This number is high compared to within-population studies of wild animals (e.g. 2.99% heritability for fitness;, *44*), indicating that hybrid incompatibilities increase mortality substantially in the F_2_ generation in *L. sinapis*. We also found that individuals that died during the larval or pupal stages had slower developmental rates (Random slopes model; *p ≈* 0.002; Figure 1D and Tables S3-4).

**Figure 1.**
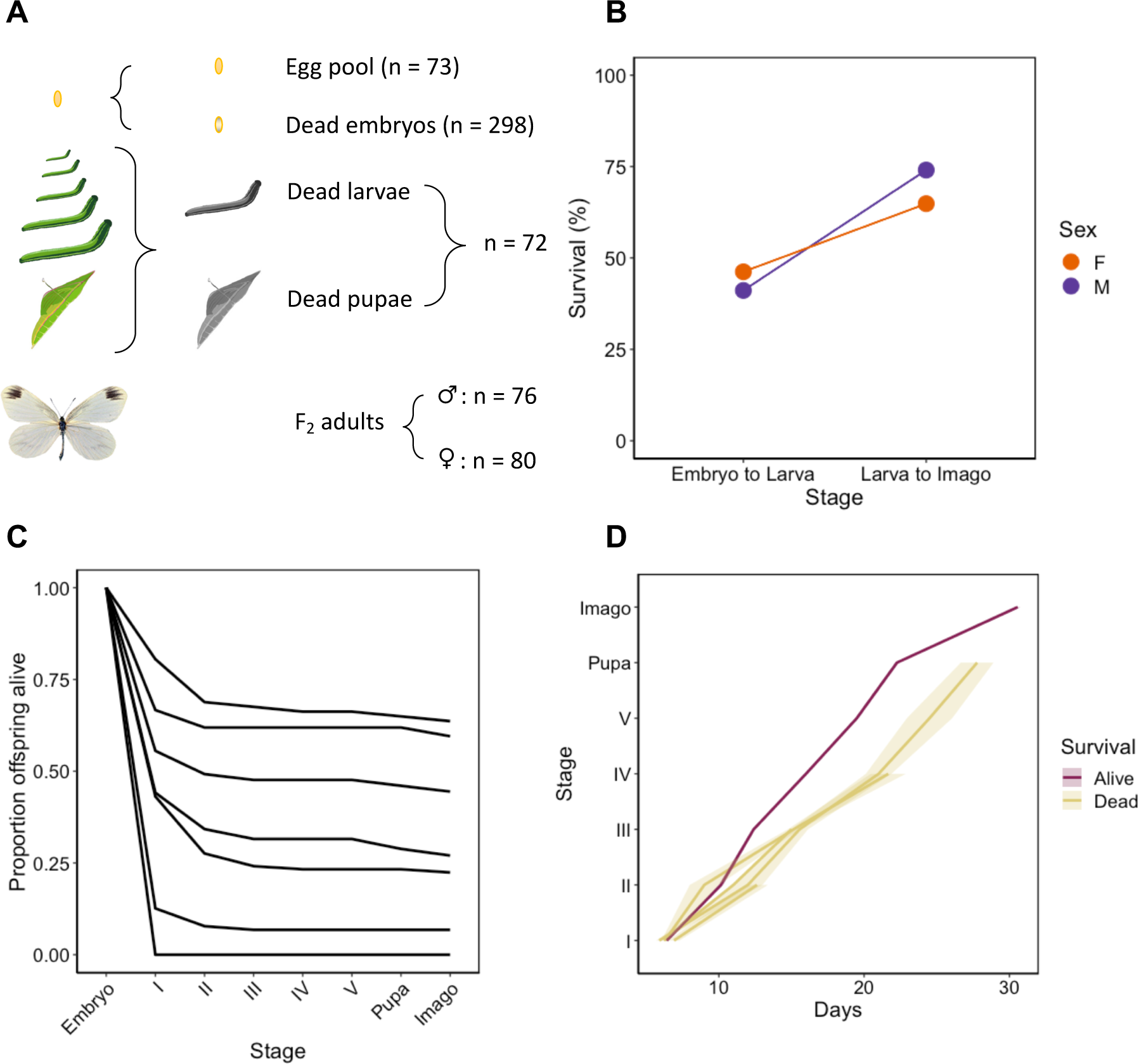
Summary of results from the F2 survival experiment produced by crosses between *L. sinapis* chromosomal races. We monitored cohorts of F2 offspring until death or emergence as adults and scored developmental stage and survival status. (A) Numbers of individuals in each pool. (B) Survival proportions across the major developmental transitions from egg to larva and from larva to imago for each sex. Overall survival was 30% for both sexes. (C) Survival curves throughout development per family. Numbers I-V show the five larval instars. (D) Comparison of the average developmental time trajectories between alive and dead F2 offspring of different lifespans. Days (X-axis) represent the time until reaching the corresponding stage. Shaded regions illustrate the 95% confidence intervals.

### Genomic architecture of F_2_ hybrid inviability

To detect genomic regions involved in hybrid inviability, we sequenced several experimental pools and compared allele frequencies between F_2_ individuals surviving to adulthood (*Alive*) and individuals that died during the larval or pupal stage (*Dead*; Figure 1A and Table S5). Using previously published population resequencing data (*45*), we identified 27,720 fixed differences between the CAT and SWE *L. sinapis*. We inferred the ancestral allele for the fixed differences using two individuals each of two outgroup species *L. reali* and *L. juvernica.* Here, 21,654 of the 27,720 fixed differences could be polarized and we found that the CAT population harbored the derived allele for 67% of the variants. We used all 27,720 fixed differences as markers to track the ancestry of genomic regions in the F_2_ pools. To correct for potential reference biases, we mapped the PoolSeq reads twice to a previously available *L. sinapis* reference genome (*46*), where all fixed differences were either set as the CAT or SWE allele. Allele frequencies for each pool were calculated as an average across both mappings. We used a generalized additive model to smooth the allele frequencies along chromosomes and to identify significantly differentiated regions between pools. To identify regions potentially associated with hybrid incompatibilities (candidate regions), we compared the allele frequencies between the *Alive* and *Dead* pools. This analysis revealed that 37 genomic regions had significantly deviating allele frequencies compared to random expectations. In the *Alive* group, 22 regions had an excess of the CAT and 15 had an excess of the SWE variant, respectively (Figure 2). Regions with an excess of the CAT variant comprised 13.5% (92.2 Mb) of the genome in the *Alive* group while regions with SWE ancestry comprised 6.6% (45.4 Mb). Candidate regions varied in size from 73 kb to 14.9 Mb and sometimes spanned entire chromosomes, such as chromosome 14 and 35 (Figure 2). As a stringent complementary method to detect significant allele frequency shifts between pools, we used the QTLseqR method on the *Alive vs*. *Dead* data set (Figure 2, Figure S3). All (n = 6) except one of the QTLs detected in this analysis were located inside four of the 37 candidate regions identified in the initial scan. We classify these six regions as large-effect loci, since they are located in genomic regions with especially pronounced allele frequency differences (Figure 2).

**Figure 2.**
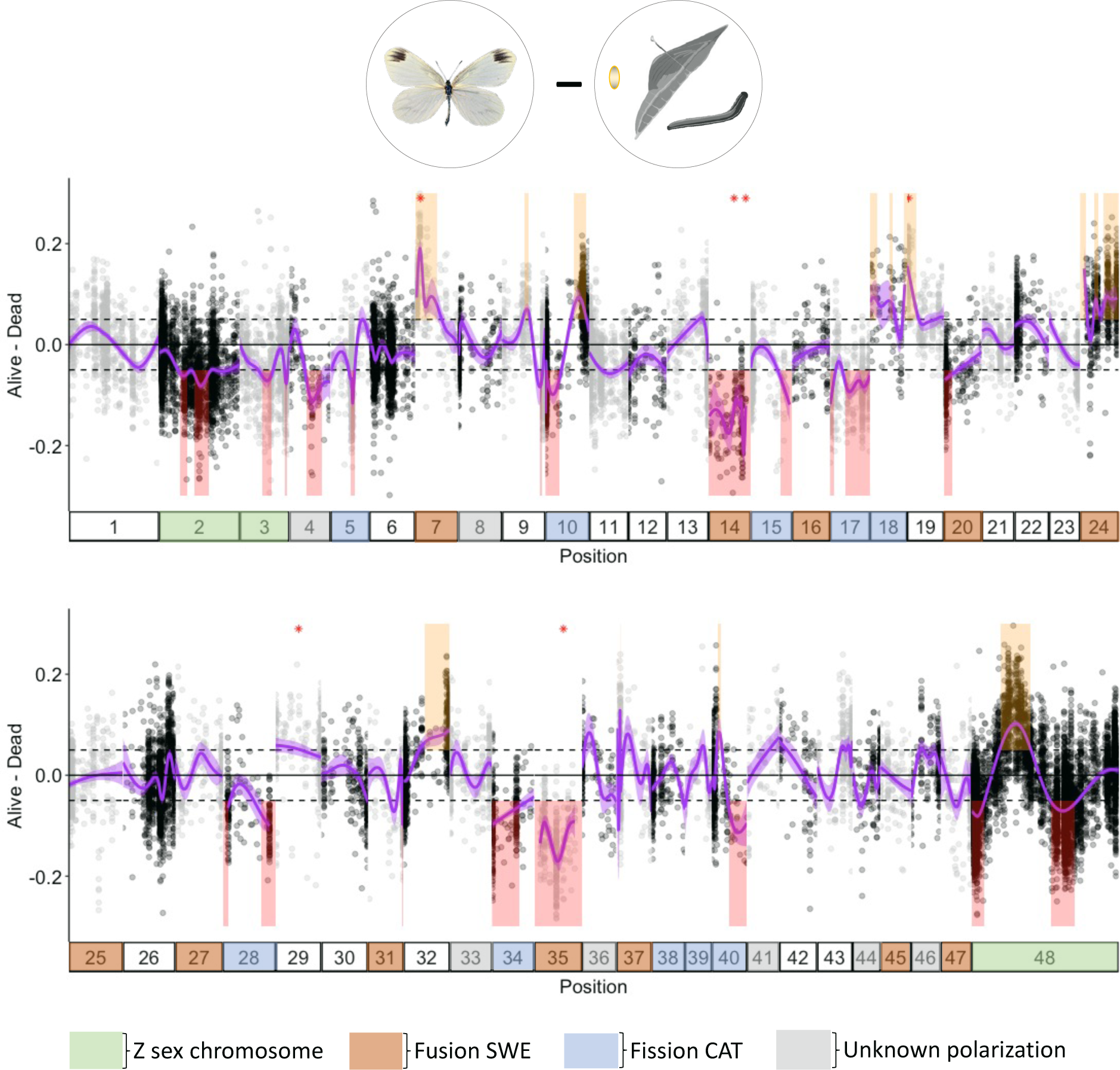
Genomic architecture of F2 hybrid inviability in *L. sinapis* mapped by comparing allele frequencies of the *Alive* and *Dead* pools. Y-axes represent the allele frequency difference between the pools *Alive* (F2 adult males and females) and *Dead* (dead embryos, larvae and pupae). X-axes show the chromosomes (numbered bars) ordered by size, except chromosome 48 which contains the ancestral Z chromosome of Lepidoptera. Dots show the position and allele frequency of the 27,240 markers, polarized for the allele frequency in the SWE population. The purple curve represents a generalized additive model fitted to the allele frequency difference between pools. Shaded areas in the graph represent regions were the *Alive* pool had an excess of SWE (yellow) or CAT (red) alleles, respectively (i.e. where the 95% CI of the curve > |0.05|). Red asterisks (*) indicate the mid position of candidate regions identified using QTLseq. Chromosomes 2, 3 and 48 (green) are the Z-chromosomes, by convention denoted Z2, Z3 and Z1 respectively. The colors of chromosomes indicate if they represent derived fusions in the SWE population (brown), derived fissions in the CAT population (blue), or segregating fission/fusion polymorphisms (grey). Note that only simple rearrangements (involving two unfused elements) are shown.

We compared allele frequencies between the *Alive* and the egg pool to test whether the candidate regions detected in the *Alive vs. Dead* comparison could be confirmed using an alternative approach. Note that this is not a strict test of repeatability given that the *Alive* pool was used in both analyses. We found that candidate regions in these comparisons overlapped 1.39-fold over the random expectation (Monte Carlo, *p* = 0.022, n = 1,000). We repeated this analysis using a more stringent (0.075) frequency difference threshold for the *Alive vs.* egg pool comparison and found similar results (odds ratio ≈ 1.76, *p* = 0.022; Figure S4). The comparison between the *Alive vs. Dead* and the *Alive vs.* egg pools was complicated by the observation that the egg pool consisted of approximately 68% females, according to the observed read coverage on Z_1_ and Z_2_, while the *Alive* and *Dead* pools had equal sex ratios (Table 1) We expect that sex-ratios of pools, within and between comparisons, affect the predicted candidate regions since the Z_1_ chromosomes are hemizygous in females which will affect the expression of incompatibilities caused by recessive variants. Consequently, a comparison between for example dead larvae and dead embryos would be confounded by the difference in sex ratios between these pools (Table 1).

**Table 1.** Inferred sex ratios of pools based on read mapping coverage of chromosome 48 (Z_1_).

| Pool | % males | Sample size |
| --- | --- | --- |
| Egg pool | 32.2% | 73 |
| Dead embryos | 50.5% | 298 |
| Dead larvae and pupae | 37.5% | 72 |

### Rearrangements and hybrid incompatibilities

Since chromosomal rearrangements might affect hybrid fitness, we investigated whether chromosomes involved in fission/fusion differences between the SWE and the CAT populations were enriched for hybrid inviability candidate regions. This analysis was performed both for the entire chromosomes involved in rearrangements in general and for the evolutionary breakpoint regions (EBRs; ± 1 Mb of an inferred fission/fusion breakpoint) more specifically. For the entire chromosomes, we found no significant enrichment of candidate regions after correcting for multiple tests (Table 2). In the EBRs, however, derived fusions were significantly enriched for candidate regions (Monte Carlo *p <* 0.02, n = 1,000). To rule out that our definition of candidate regions cause a spurious association, we also tested the six large-effect loci identified using the QTL-analysis. These loci were also significantly enriched on chromosomes involved in derived fusions (odds ratio ≈ 4.32, *p <* 0.001), but not EBRs (Table S6).

**Table 2.** Associations between chromosomal rearrangements and hybrid inviability candidate regions. The analysis was performed for the entire chromosomes, evolutionary breakpoint regions (EBRs) and non-EBR ends of chromosomes, respectively. Chromosomes with unknown polarization are those with fission/fusion polymorphisms that are known to segregate in different *Leptidea* species.

| Category | Polarization | Odds ratio | *p*-value | *p*-value* |
| --- | --- | --- | --- | --- |
| Chromosome | Fission CAT | 1.768 | 0.034 | 0.102 |
| Chromosome | Fusion SWE | 1.543 | 0.076 | 0.228 |
| Chromosome | Unknown | 0.312 | 0.272 | 0.816 |
| EBR | Fission CAT | 0.996 | 0.110 | 0.330 |
| EBR | Fusion SWE | 1.992 | 0.002 | 0.006 |
| EBR | Unknown | 1.245 | 0.223 | 0.672 |
| non-EBR ends | Fission CAT | 2.619 | < 0.001 | < 0.001 |
| non-EBR ends | Fusion SWE | 1.992 | < 0.001 | < 0.001 |
| non-EBR ends | Unknown | 0 | < 0.001 | < 0.001 |

\*Corrected for multiple testing using the Bonferroni method for each category separately.

The association between fusion EBRs and candidate regions could be due to an association between chromosome ends and hybrid inviability, rather than the fusion event itself. Consequently, we also investigated non-EBR ends of rearranged chromosomes. This analysis showed that there was an enrichment of candidate regions within derived fusions (odds ratio ≈ 1.99; *p <* 0.001) and fissions (odds ratio ≈ 2.62; *p <* 0.001). Chromosomes with unknown polarization, however, contained no candidate regions at non-EBR ends (odds ratio = 0; Table 2). To further assess if the association between candidate regions and EBRs could be a consequence of differences in gene density between conserved and rearranged regions, we investigated the relationship between coding sequence (CDS) density and the candidate regions. This analysis unveiled a small but significant excess (odds ratio ≈ 1.08; *p* = 0.012) of CDS regions in candidate regions compared to the genome-wide level (Table S7). Derived fusion EBRs had a significantly lower density of CDS regions compared to the genome-wide level (odds ratio ≈ 0.58; *p* = 0.008). Thus, CDS density cannot explain the association between candidate regions and fusion EBRs.

So far we have only considered simple chromosomal rearrangements, i.e. fissions in the CAT population or fusions in the SWE population, resulting in a 2:1 homologous chromosome number ratio for the CAT:SWE population pair. Previous analyses have shown that multiple complex chromosome chain rearrangements are segregating within and between the SWE and CAT *L. sinapis* populations (*36*). In addition, *Brenthis* butterflies show reduced gene flow at complex rearrangements (*47*). We therefore assessed if chromosomes involved in chain rearrangements (chromosomes 6, 13, 21, 22, 26, 30) were enriched in candidate regions. This analysis showed chain rearrangements had significantly fewer candidate regions than expected by chance (*p* < 0.001).

Chromosomal inversions are prime examples of rearrangements that can reduce the crossover rate of heterokaryotypes. We characterized inversions between the two populations with whole-genome alignments of chromosome-level assemblies of CAT and SWE males. The analysis revealed 20 inverted regions between the SWE and the CAT reference. The length of the inversions ranged from 12.6 to 616.4 kb and five of the inversions intersected with hybrid inviability candidate regions. This was 1.5-fold higher than the random expectation, but not statistically significant (Monte Carlo p = 0.552; n = 1,000).

### No indications of systematic aneuploidy in dead embryos

Chromosome fission/fusion polymorphisms can lead to non-disjunction during meiosis and formation of aneuploid gametes (reviewed in *9*). In some cases, aneuploid gametes can survive many rounds of cell-division (*16*). Investigating aneuploidy can therefore inform about the mechanisms relating chromosome fusions and hybrid inviability. If non-disjunction during meiosis due to fusions causes hybrid inviability we expect to see systematic aneuploidy. If there is no systematic aneuploidy, we expect the relationship between fusions and hybrid inviability to be indirect and in that case hybrid inviability is more likely a consequence of linked genic incompatibility factors. We investigated whether any systematic aneuploidies were present in the dead embryo pool by comparing read coverage among chromosomes at fixed differences. The autosome with the highest coverage had 37% higher coverage than the average level among all autosomes (Figure S5). In the case of aneuploidy, we would expect single autosomes to have either 50% higher coverage (trisomy), half the coverage (monosomy) or no coverage at all (nullisomy), compared to other autosomes. For surviving F_2_ males and females, the differences between the highest covered autosome and the average were 27% and 35% respectively. For both dead embryos and adult survivors, chromosomes 17 and 21 had the highest coverage (Table S8). Neither of these two chromosomes is associated with simple derived fusions (see Figure 2). This indicates that it is unlikely that systematic aneuploidies are present in the dead embryo pool and that the relationship between hybrid inviability and fusions is caused by other factors. Consequently, we further examined the indirect mechanism of chromosomal speciation by investigating the relationships between hybrid inviability, chromosome fusions and the recombination rate.

### Hybrid inviability candidate regions are characterized by low recombination rates

To test if hybrid inviability candidate regions show a reduced recombination rate compared to other parts of the genome, we bootstrapped genomic regions of the same sizes as the observed candidate regions and extracted observed recombination rates in those regions from population-specific linkage maps (Figure 3 and Figure S6). Importantly, since the underlying regional recombination rate variation can affect the size distribution of potential regions with restricted gene flow, we calculated the arithmetic mean recombination rate without normalizing for sequence length. We found that the recombination rates in candidate regions in both the SWE (2.44 cM/Mb; Monte Carlo *p* ≈ 0.046, n = 100,000) and the CAT population (3.32 cM/Mb; *p ≈* 0.014) were significantly lower compared to the genome-wide rates (3.03 and 4.25 cM/Mb for SWE and CAT respectively; Figure 3).

**Figure 3.**
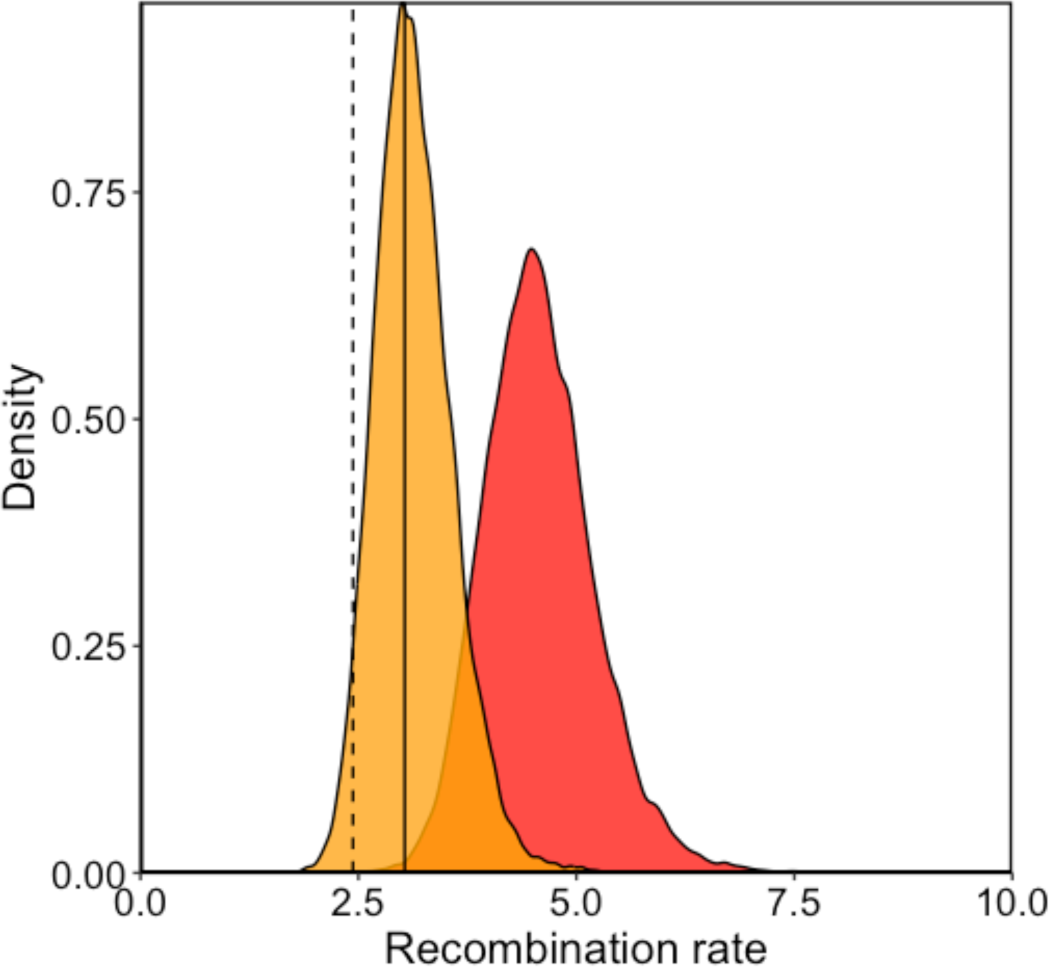
Parental population recombination rates in candidate regions for hybrid inviability. Distributions of the genome-wide recombination rates determined by resampling for the SWE (orange) and the CAT (red) *L. sinapis* populations. The vertical solid and dashed lines show the observed average recombination rates of the candidate regions for the CAT and the SWE population, respectively.

### Fusions are associated with low recombination rates in both arrangements

Candidate regions for hybrid inviability were generally clustered in regions with reduced recombination rates. Low recombination rates could be the explanatory factor of the association between hybrid inviability and chromosome fusions. When a chromosome fusion occurs, loci in the vicinity of the fusion point that were segregating becomes tightly linked. Low recombination rates near fusions is expected to extend over a larger area since the center of large chromosomes in butterflies tend to show reduced recombination rates compared to the genome average (*33*, *48*). In line with this we found that derived fusion EBR regions (± 1 Mb of an inferred breakpoint), had significantly reduced recombination rates compared to the genome-wide rate in both the SWE (fused state) population (one sample Wilcoxon tests; *p* < 0.05; Figure 4 and Table S9). In addition, the CAT (unfused state) had also lower recombination rates (*p* < 0.05; Figure 4 and Table S9), in line with low recombination rates at chromosome ends in Lepidoptera (*33*, *48*). EBR and non-EBR ends of fusions did not have significantly different recombination rates (*p >* 0.05). We also investigated the derived fissions and fission/fusion polymorphism with unknown polarization and found low recombination rates compared to genome-wide rates at non-EBR ends but not EBRs (Figure S7 and Table S10). In conclusion, both fused and unfused chromosomes had reduced recombination rate in the EBRs for derived fusions.

**Figure 4.**
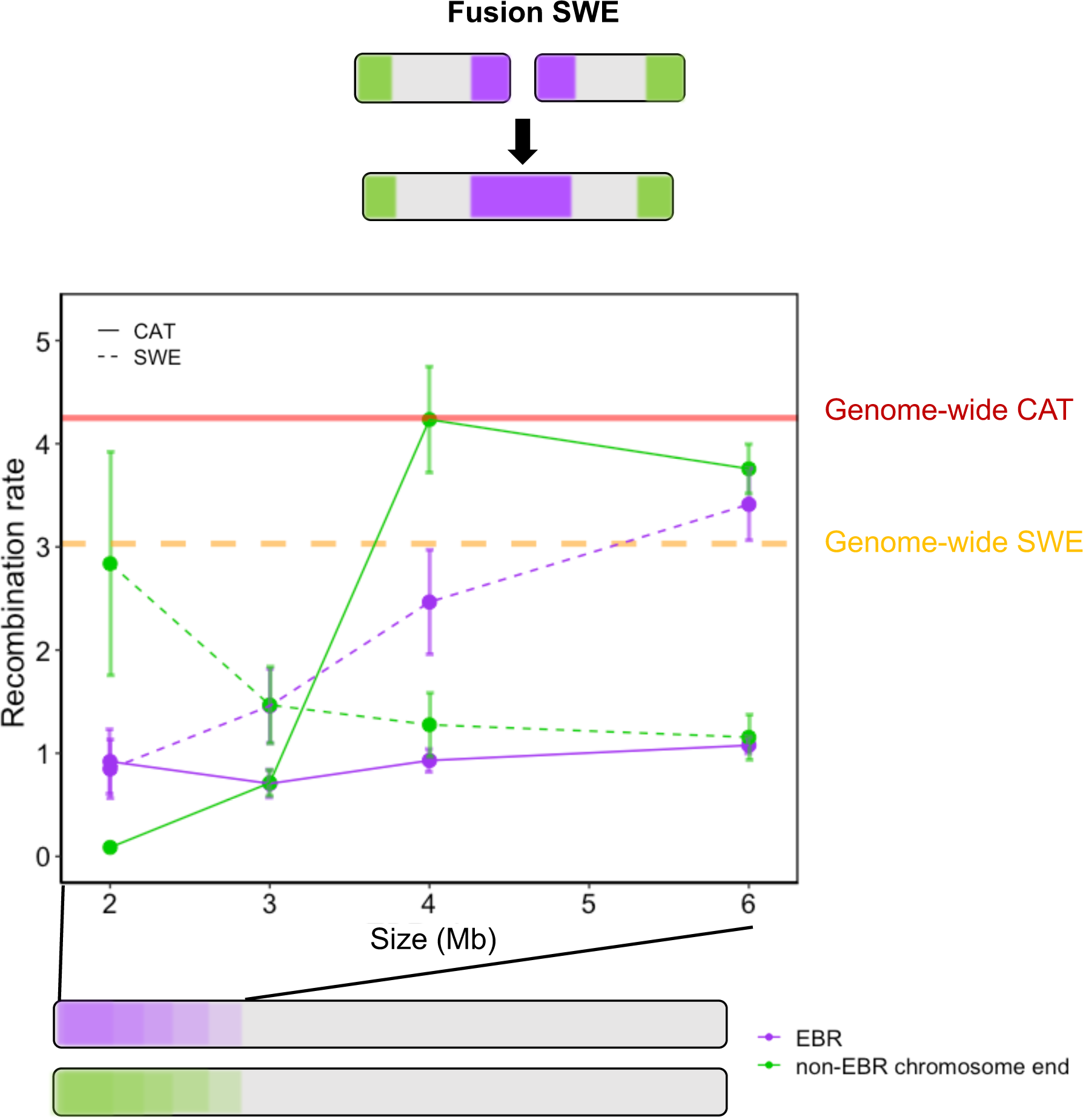
Patterns of recombination near chromosome fusion evolutionary breakpoint regions (EBRs). EBRs are shown in purple and non-EBR chromosome ends are shown in green. Patterns of average parental recombination rates in EBRs and non-EBRs chromosome ends are presented for 2, 3, 4 and 6 Mb windows. Error bars represent the standard error of the mean. Solid and dashed lines show the recombination rates in the CAT and the SWE population, respectively. Horizontal lines represent mean genome-wide recombination rates for the CAT (red) and SWE (orange) population.

### Low levels of gene flow during divergence

To further understand the evolutionary origins of hybrid inviability, we investigated the demographic history and genomic landscape of differentiation and divergence using population resequencing data from 10 males each of CAT and SWE *L. sinapis*. First, we investigated the demographic history of the populations using GADMA (Figure 5). Models incorporating gene flow provided a superior fit to the observed joint minor allele frequency spectrum compared to models without migration (d_AIC_ = 3,715; Figure 5 and Table S11). Inferred gene flow was low in general, but higher from the SWE to the CAT population (M_SWE_è_CAT_ = 1.07, 95% CI: 0.5–1.6) than vice versa (M_CAT_è_SWE_ = 0.18, 95% CI: 0 – 0.4; Figure 5 and see Table S11 for inferred parameter values). The low level of gene flow was reflected in the genomic landscape of differentiation, estimated using *F_ST_* which compares heterozygosity within and between populations. Average *F_ST_* in non-overlapping 10 kb windows across the entire genome was 0.26. We also computed the level of absolute differentiation (*D_XY_*) between the populations and level of genetic diversity (π) within each population. The genome-wide average *D_XY_* was 0.012, slightly higher than the population specific estimates of diversity (π_SWE_ = 0.0085 and π_CAT_ = 0.0093). A positive association between *F_ST_* and *D_XY_* across the genome can be a signature of regional variation in resistance to gene flow between incipient species (*49*, *50*). We therefore compared the window-based estimates of *F_ST_* and *D_XY_* and found a weak but significant positive correlation (Spearman’s ρ = 0.11; *p <* 2.2 *10^-16^; Figure S8), indicating a minor impact of gene flow on the genomic landscape of differentiation between SWE and CAT *L. sinapis*.

**Figure 5.**
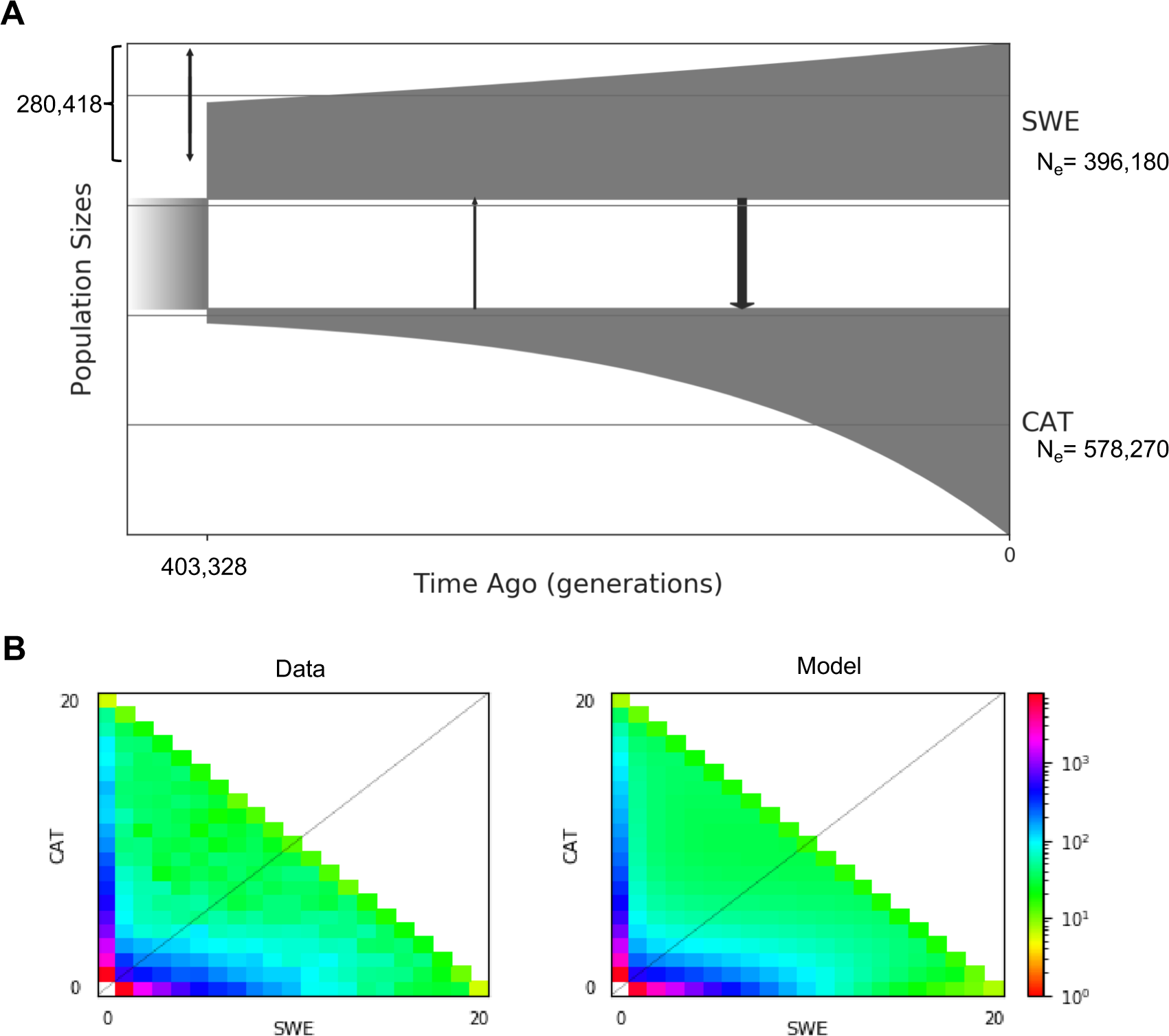
Demographic history of SWE and CAT *L. sinapis* inferred from population resequencing data. (A) Schematic model of the inferred history. Sizes of boxes represent the effective population size (*Ne*). Arrows connecting boxes are directional migration rates averaged across the entire epoch. Arrow widths are scaled to illustrate the intensity of migration. (B) Computed joint minor allele frequency spectrum (left) and predicted minor allele frequency spectrum from the model (right).

### Higher levels of genetic differentiation in hybrid inviability candidate regions

We contrasted population genetic summary statistics in candidate regions with the rest of the genome to get a better understanding of the processes that may have influenced their evolution. On average, *F_ST_* between parental samples (CAT and SWE) measured in 10 kb windows was slightly higher in candidate regions (0.2728) compared to non-candidate regions (0.2602, Table 3). To control for chromosome effects, such as a higher expected differentiation of the Z chromosomes (Figure S9), we performed an analysis of variance (ANOVA) using candidate region status and chromosome identity as fixed effects. This analysis revealed a significantly higher *F_ST_* in candidate regions than in non-candidate regions (Table 4, Figure 6A). Conversely, π_CAT_ was significantly lower in candidate regions than in non-candidate regions. We found no significant differences in *D_XY_* and π_SWE_ in candidate regions compared to the rest of the genome when controlling for between-chromosome variation. We also tested models including coding sequence (CDS) density as a predictor, with qualitatively similar results (not shown). To summarize, most hybrid inviability candidate regions (but not all, see Figure 6B and Figure S10) showed elevated genetic differentiation and reduced π_CAT_.

**Table 3.** Estimated average [95% confindence intervals] population genetic summary statistics in non-overlapping 10 kb windows for candidate and non-candidate incompatibility regions in the genome, respectively.

| Statistic | Candidate regions | Non-candidate regions |
| --- | --- | --- |
| $F_{ST}$ | 0.2728 [0.2704 – 0.2751] | 0.2602 [0.2591 – 0.2613] |
| $D_{XY}$ | 0.0115 [0.0104 – 0.0126] | 0.0122 [0.0121 – 0.0122] |
| $\pi_{SWE}$ | 0.0081 [0.0081 – 0.0082] | 0.0086 [0.0086 – 0.0087] |
| $\pi_{CAT}$ | 0.0087 [0.0087 – 0.0088] | 0.0094 [0.0093 – 0.0094] |

$F_{ST}$ , $D_{XY}$ and $\pi$ were rounded to four decimals.

**Table 4.** Results from the ANOVA analysis of differences between population genetic summary statistics (estimated in 10 kb windows) inside and outside candidate regions for hybrid incompatibility between SWE and CAT *L. sinapis*. Chromosome and Status (within or outside candidate regions) represent the fixed effect predictors.

| Response variable | Predictor variable | $F$ | $p$ |
| --- | --- | --- | --- |
| $F_{ST}$ | Chromosome | 132.322 | $< 2.2 \cdot 10^{-16}$ |
| | Status | 37.393 | $9.709 \cdot 10^{-10}$ |
| $D_{XY}$ | Chromosome | 106.011 | $< 2.2 \cdot 10^{-16}$ |
|  | Status | 0.601 | 0.438 |
| $\pi_{SWE}$ | Chromosome | 146.032 | $< 2.2 \cdot 10^{-16}$ |
|  | Status | 0.380 | 0.538 |
| $\pi_{CAT}$ | Chromosome | 77.812 | $< 2.2 \cdot 10^{-16}$ |
| | Status | 19.918 | $8.095 \cdot 10^{-6}$ |

$F$ and $p$ were rounded to three decimals.

**Figure 6.**
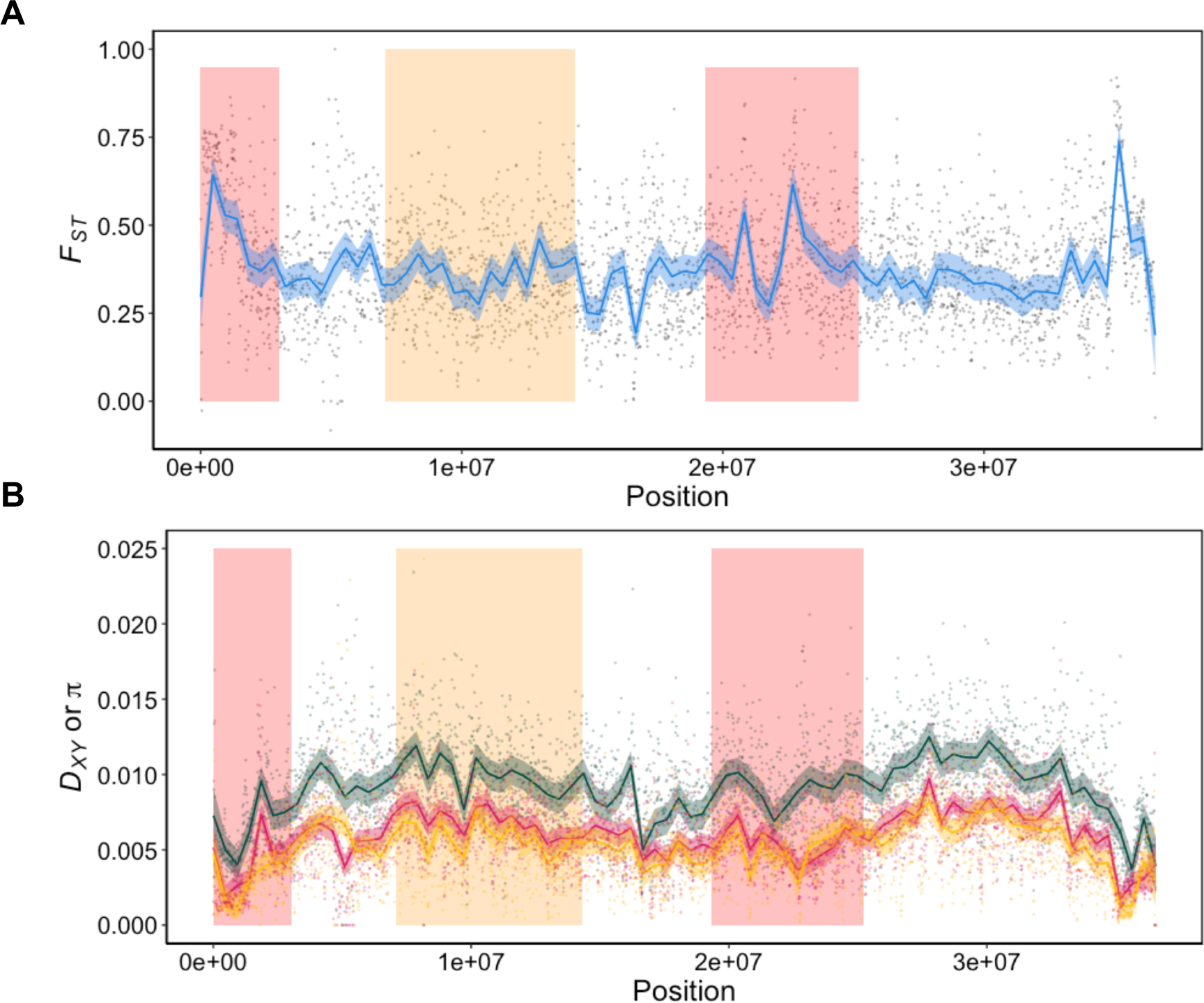
Population genetic summary statistics across chromosome 48 (Z1). (A) 10 kb window-based estimates of genetic differentiation (*FST*). (B) Absolute divergence (*DXY,* dark green line), and genetic diversity in the CAT (πCAT, red line) and SWE (πSWE, orange line) population of *L. sinapis*, respectively. Values have been smoothed using local regression and shaded regions represent 95% confidence intervals. Boxes represent candidate regions for hybrid incompatibility between SWE and CAT *L. sinapis*. This example chromosome shows that in some candidate regions there are peaks of *FST* (region 1 and 3 from left to right). In others, there are no clear peaks in *FST* (region 2). There were also regions with elevated *FST* such as the 3’ region of chromosome 48, which were not associated with hybrid inviability.

## Discussion

Here we used a combination of approaches to investigate the genetic underpinnings of hybrid inviability between two populations of wood whites that differ in karyotype structure due to a large number of chromosomal rearrangements. Our detailed characterization of survival in hybrid offspring, the mapping of candidate hybrid inviability loci and investigations of the differences in population genomic signatures at candidate loci compared to the genome in general revealed that the evolution of hybrid inviability is associated with the extensive chromosomal rearrangements that have occurred in different lineages of *L. sinapis*.

### The genetic basis of hybrid inviability - comparisons of dead and alive offspring

Whole-genome sequencing of individuals is the ideal method for characterizing the genetic basis of hybrid incompatibility, since such data allow for assessment of genetic linkage between loci, which is key for uncovering the epistatic relationships that are integral to the BDMI model (*51*). Nevertheless, sequencing of pooled samples (PoolSeq) is an alternative strategy that allows mapping in hundreds or thousands of individuals at a reasonable cost (*52*). Here we added a novel twist to the PoolSeq approach of detecting hybrid inviability in a natural system by sampling both surviving and dead F_2_ offspring from crosses between parental lineages with distinct karyotypes.

To get a detailed understanding of the core genetic underpinnings of a speciation event, we would ideally target the mechanisms that lead to reproductive isolation at the onset of the speciation process (*1*). For the genetic mapping of reproductive isolation this poses a dilemma, since the level of divergence between incipient species pairs is positively associated with the number of markers available for mapping (*43*, *53*). Here we demonstrated that it is possible to both identify hybrid inviability candidate regions and extract informative data about the evolution of hybrid inviability in a cross of a non-model organism, using a limited set (∼ 27,000) of informative genetic markers. It is important to emphasize that the identified loci are no more than candidate regions and further experiments would be necessary to identify the genes and specific genetic differences involved, as well as the potential epistatic interactions that cause hybrid inviability. We further outline three of the challenges with the PoolSeq method to study hybrid incompatibilities below. First, since we expect recessive hybrid incompatibilities to be at play in a system with F_2_ hybrid breakdown, such as in *L. sinapis*, each mating will generate a relatively high proportion of viable offspring and allele frequency deviations from the expected value of 0.5 will in most cases be modest (*43*). This reduces the power to detect loci associated with inviability. Here we compared allele frequencies between pools of both *Alive* and *Dead* F_2_ offspring which increases this power to some extent. This is because alleles enriched for the variant with SWE ancestry in the *Alive* pool necessarily will be enriched for the variant with CAT ancestry in the *Dead* pool, and vice versa. Second, a number of deaths from a cross can be due to environmental effects associated with lab conditions. While our data point to a relatively strong genetic component for survival (*h^2^ =* 0.38), such environmental deaths will decrease the power for identification of hybrid inviability loci, thereby making the approach somewhat conservative. Third, if hybrid inviability is caused by a combination of (a few) large effect and (many) small effect loci (i.e. the trait is polygenic), as have been observed for example for hybrid male sterility in the *Drosophila simulans* clade (*54*), the power to detect the true genomic architecture in an F_2_ cross with a limited number of recombination events will be low. For the present study it means that we cannot exclude that there are many more loci with small effects involved in hybrid inviability. We did however detect a set of candidate regions with a sufficiently large effect sizes which allowed for further investigation of the evolution of hybrid inviability.

### Genomic architecture of hybrid inviability is concentrated to low-recombining fast evolving regions of the genome

In theory, hybrid incompatibilities could evolve in any region of the genome where novel mutations or sorting of ancestral variants differ between diverging lineages. However, incompatibilities are more likely to fix in regions with faster substitution rate/lineage sorting, such as regions with low recombination rates. In agreement with this prediction, we observed that hybrid inviability candidate regions had higher *F_ST_* and significantly lower recombination rate compared to non-candidate regions. Sex chromosomes often diverge at faster rates than autosomes and are often highlighted as hotspots of hybrid incompatibilities (“Haldane’s rule” and the “Large X/Z-effect”) (*55–57*). However, we observed equal survival of males and females, supporting that the role of sex chromosomes is more important for the fitness (in particular sterility) of F_1_ hybrids than F_2_ hybrids (*1*). In addition, none of the large-effect loci identified mapped to the Z chromosomes. Instead, we observed an enrichment of hybrid inviability candidate regions at the ends of chromosomes that have been involved in chromosomal rearrangements between SWE and CAT *L. sinapis.* Some of these EBRs between karyotypes are characterized by low recombination rates. This is not entirely surprising. Theoretical work on speciation has emphasized the reduction of recombination as a mechanism promoting reproductive isolation (*22*, *23*, *25*, *31*, *58*), and hybrid incompatibilities have been mapped to low-recombination regions in several systems (*17*, *59*). In most cases this has involved inversions, which only show reduced recombination rate in heterokaryotypes. A low recombination rate in certain genomic regions in homokaryotypes could potentially also be associated with a lower recombination rate in the homologous regions in heterokaryotypes. We currently lack data about recombination rate variation in heterokaryotypes, which would be needed to truly compare the effects. However, our results show that hybrid inviability candidate regions are enriched at chromosome fusion breakpoints, but not in inversions, which presumably only have reduced recombination rate in heterokaryotypes. This suggest that reduced homokaryotype recombination rate associated with chromosome fusions in some cases can be more important than recombination restriction in heterokaryotypes alone.

### Does recombination fully explain the association between chromosome fusions and hybrid inviability?

We observed low recombination rates in both candidate regions for hybrid incompatibility and in fusion EBRs. This raises the question whether the association between chromosome fusions and hybrid inviability is fully mediated by the effect of chromosome fusions on recombination? Since we also observed a significant association between hybrid inviability candidate loci at non-EBR ends of derived fusions it is possible that recombination is the underlying factor explaining the association between chromosome fusions and hybrid inviability. However, we did not see a significant enrichment of hybrid inviability candidates in the low recombination non-EBRs for chromosomes segregating among the most closely related *Leptidea* species. This means that more recently evolved rearrangements, such as the fusions in the SWE lineage are more likely to be associated with hybrid inviability. This indicates that low recombination rate by itself may not fully explain the association between fusions and hybrid inviability. Instead, *when* a rearrangement occurred in the evolutionary history appears to be important for whether it harbors hybrid inviability factors or not. However, if recent fixation of rearrangements were the sole explanation for the evolution of hybrid inviability we would expect fission EBRs to be enriched for incompatibility loci as well. This was not the case, which indicates that it is both the low recombination rate associated with fusion breakpoints and the recent fixation of fusions that mediates the association with the evolution of hybrid inviability. Derived fusions may have fixed due to selection for increased linkage disequilibrium between alleles at loci located near the ends of two unfused chromosomes (*31*, *60*). Importantly, loci driving the fixation of fusions could be but are not necessarily the same loci causing hybrid inviability.

### Evolution of hybrid inviability through association with chromosome fusions

A genetic correlation between traits can either be caused by pleiotropy, tight physical linkage between independent genes affecting the traits, or both. In the case of hybrid inviability and chromosome fusions, a pleiotropic mechanism would be that hybrid inviability is caused directly by the changed chromosome structure itself. This could for example be caused by aneuploidies arising from non-disjunction in F_1_ meiosis or mitotic mis-segregation in the embryo (*16*). A physical linkage explanation would instead be that the fixation of a chromosomal rearrangement leads to the fixation of linked genic incompatibility factors. Both pleiotropy and physical linkage could be cooccurring, as has been shown in F_2_ crosses among populations of the Australian grasshopper *Caledia captiva* differing either in multiple rearrangements, fixed genetic differences or both (*61*). Since we observed no systematic aneuploidies, our data supports the physical linkage model, i.e., an indirect relationship between chromosome fusions and hybrid inviability. The physical linkage model has been empirically supported in monkeyflowers (*Mimulus guttatus*) where hybrid inviability has evolved between copper-tolerant and non-copper-tolerant populations and a gene involved in adaption to copper-polluted soil has been shown to be tightly linked to another gene that underlies hybrid inviability (*62*). We observed a significantly reduced recombination rate in 2 Mb regions flanking fusion points, which increases the level of linkage disequilibrium leading to the indirect evolution of hybrid inviability. The next step would be to investigate whether fusions fixed by selection, drift, or a fixation bias (*63*). If fusions fixed by selection for local adaptation, then the situation in *Leptidea* would be similar to *Mimulus*.

### Ongoing speciation or speciation reversal?

The CAT and the SWE populations of *L. sinapis* represent the most extreme cases of intraspecific karyotype variation of any diploid animal. This striking variation in karyotype setup is further characterized by a chromosome number cline, where populations in the south-western part of the distribution range have the highest and populations in the northern (e.g. SWE) and eastern (e.g. Kazakhstan, KAZ) parts the lowest number of chromosomes. KAZ and SWE populations also have low genetic differentiation (*34*, *45*). The SWE and CAT populations therefore most likely represent an eastern and a western ancestry group, respectively. According to our demographic analysis they cannot have shared a refugium during the last-glacial maximum (∼ 20 kya) and accumulated genetic differences thereafter, which has previously been proposed (*34*). The remarkable chromosome number differences between *L. sinapis* populations have also been used to argue for a clinal speciation model (*12*, *34*). However, the relatively deep divergence time between the SWE and the CAT population indicates that the current chromosome number cline is a consequence of secondary contact and that we might be witnessing a case of ‘speciation reversal’ in this system. The inferred historical gene flow was low between these ancestry groups and absolute divergence, *D_XY_,* was not elevated in candidate regions. This indicates that both chromosome number differences and hybrid inviability evolved during the repeated Pleistocene glaciations, when an eastern (represented by SWE) and a western (represented by CAT) group of *L. sinapis* were isolated from each other. These refugial populations probably came into secondary contact because of post-glacial population expansions. Hence, populations throughout central Europe and the British Isles likely have ‘hybrid ancestry’ and constitute a transition zone where gene-flow has occurred between ancestry groups. More detailed biogeographical analyses of *L. sinapis* in general and the central European populations in particular will be needed to verify the suggested hypothesis. Quantification of hybrid fitness in crosses between a central European population and the SWE and the CAT population, respectively, would for example be informative for understanding patterns of postzygotic isolation and potential associations between hybrid fitness, ecology and chromosomal rearrangements. Such efforts are key for the field of speciation genetics in general, since our knowledge about reversal of intrinsic postzygotic isolation is limited (*64*).

## Materials and methods

### Study system and crosses

The wood white (*Leptidea sinapis*) is one of around a dozen Eurasian *Leptidea* species which belong to the Dismorphinae subfamily (family *Pieridae*) and it has the most extreme intraspecific diploid karyotype variation of any eukaryote (*34*). The diploid chromosome number (2n) ranges from 57, 58 in SWE to 106 - 108 in CAT (*35*). Previous analyses have shown that hybrids generated by crossing the most extreme karyotypes express hybrid breakdown from the F_2_ generation and onwards (*35*). We therefore crossed SWE and CAT *L. sinapis* to establish a large set of F_2_ individuals that we could use to characterize the genetic underpinnings of hybrid inviability. Two ♀SWE x ♂CAT and five ♀CAT x ♂SWE (all offspring of wild-caught individuals) pairs were crossed in the lab in 2018. F_1_ offspring were diapaused at 8°C in a cold room and eight F_1_ x F_1_ crosses were performed in the spring of 2019. Mated F_1_ females were separated in individual jars where they had access to sugar water and bird’s-foot trefoil (*Lotus corniculatus*) for egg-laying. Females were transferred to new jars with fresh host plants and sugar water every day until they stopped laying eggs. A maximum of 10 of the first-laid eggs from each female (n = 10 from seven females and n = 3 from one female; n = 73 in total) were sampled three days after laying (the ‘egg pool’). F_2_ offspring were reared in individual jars with *ad libitum* access to the host plant *L. corniculatus*. All jars were kept in a room that varied in temperature between 23-27°C under a 16:8 hours (h) light:dark regime until 28/5 2019, and a 20:4 h regime thereafter.

### Survival experiment

All egg-laying jars were monitored daily for hatched F_2_ offspring (n = 530). After hatching, Instar I, larvae were separated into individual jars with access to *ad libitum L. corniculatus*. All individual F_2_ offspring were monitored and developmental stage (and time of day) were scored daily until they were found dead or emerged from the chrysalis as imagos. Individuals that were found dead were immediately stored at -20°C. We classified embryos as dead if they had not emerged from the egg after 9 days. Emerged imagos (the ‘*Alive*’ category) were sacrificed and stored in -20°C.

### DNA extractions and sequencing of pools

DNA was extracted using standard phenol-chloroform extraction protocols. DNA from dead larvae, pupae and imagines was extracted for each individual separately while eggs were extracted in pools of 2-21 individuals, grouped by dam. Illumina TruSeq PCR-free library preparations and whole-genome re-sequencing (2×151 bp paired-end reads with 350 bp inserts) on one Illumina NovaSeq6000 (S4 flowcell) lane were performed by NGI, SciLifeLab, Stockholm.

### Population resequencing data, variant calling, and inference of fixed differences

To track ancestry of alleles in the F_2_ offspring pools, we inferred fixed differences using individual whole-genome population re-sequencing data from 10 CAT and 10 SWE *L. sinapis* males, as well as two *L. juvernica* and two *L. reali* males (*45*). Reads < 30 bp long and with a Phred score < 33 were removed and adapters were trimmed using TrimGalore ver. 0.6.1, a wrapper for Cutadapt ver. 3.1(*65*). Trimmed reads were mapped to the Darwin Tree of Life reference genome assembly for *L. sinapis* – a male individual from Asturias in northwestern Spain with karyotype 2n = 96 (*46*) – using bwa *mem* ver. 0.7.17 (*66*). Variants were called with GATK (*67*), quality filtered with standard settings (Table S12) and used as a training set for base-quality score recalibration (*68*). Recalibrated reads were subsequently used for a second round of variant calling. Fixed differences were inferred as SNPs (i.e., excluding indels) with different alleles present in all 10 CAT and SWE individuals, respectively, allowing no missing data (n = 27,720). Ancestral state was inferred using parsimony with the requirement that at least four outgroup (i.e., *L. juvernica* and/or *L. reali*) chromosomes harbored a specific allele of the inferred fixed variants (n = 21,654).

### Pool-seq read mapping and variant calling

Pool-seq Illumina paired-end reads were trimmed and adapters were removed using TrimGalore ver. 0.6.1, a wrapper for Cutadapt ver. 3.1(*65*). In addition, seven bp were trimmed from the 3’ end of all reads with a Phred score < 33. Quality-filtered reads were aligned to two modified versions of the Asturian *L. sinapis* reference genome assembly (*46*), using bwa *mem* ver. 0.7.17(*66*). To reduce the impact of potential reference bias, we repainted the reference prior to mapping using either the CAT or the SWE allele for all inferred fixed differences, i.e. a ‘Swedenized’ and ‘Catalanized’ reference, respectively. For downstream analysis, we used the average allele frequency for both mappings. Deduplication was performed using Picard *MarkDuplicates* ver. 2.23.4 and reads with mapping quality < 20 were removed. Allele frequencies were estimated for fixed variants using MAPGD *pool* (*69*). Only markers with a likelihood ratio *p* < 10^-6^ were kept for downstream analysis. Allele frequencies used in downstream analyses were polarized for SWE ancestry.

### Quantitative genetics analyses

We tracked the pedigree of all F_2_ offspring in the survival experiment and performed a quantitative genetic analysis to determine the heritability for survival. Genetic variance-covariance matrices were computed using the R package Nadiv (*70*). Heritability was determined using Bayesian inference of the “animal model” as implemented in the R package mcmcGLMM (*71*, *72*). In this framework, posterior values of genetic variance are sampled from a prior distribution and parameter space is explore using Markov Chain Monte Carlo methods to form a posterior distribution of genetic variance. In the first model, we used survival as the response variable and the genetic variance-covariance matrix as the random predictor to quantify the narrow-sense heritability in survival. Since survival is a binary trait, we used a threshold link function. Both the uninformative prior (*V* = 1, *nu* = 1^-6^) and a parameter-expanded prior (*V* = 1, *nu* = 1, *alpha.mu* = 0, *alpha.V* =1000) were applied. Both prior settings resulted in an estimated heritability within one percentage point of each other, indicating low influence of the prior settings on the posterior distribution (Table S1-2 and Figure S2). To calculate the heritability on the observed data scale, we used *model=binom1.probit* in the R package QGglmm (*73*). For models with development time as a Gaussian response variable, we used random slopes (*random = ∼ us(1+Stage):animal+animal*) and *Sex*+*Survival* as a fixed effect. We used both parameter-expanded priors and uninformative priors and both settings gave qualitatively similar results (Tables S3-4).

### Inference of candidate regions

Candidate regions for hybrid inviability were characterized by calculating allele frequency differences between the *Alive* (adult males and females) and the *Dead* (dead embryos and dead larvae + dead pupae) pools. We used all 27,240 markers with data for all 2 × 4 sequence pools x mapping combinations. A generalized additive model (*y ∼ s[x, bs = “cs”]*) was used to get allele frequency trajectories along each chromosome. Candidate regions were defined as regions where the 95% confidence interval exceeded an absolute allele frequency difference cutoff (in general 5%, but we also applied stricter cutoffs for comparison, see below). The allele frequency differences between genomic regions are expected to be small on average, since most haplotypes are expected to be fit for a typical recessive two-locus incompatibility (*43*, *74*, *75*). As an alternative method, we performed a bulk-segregant QTLseq analysis (*76*), using the R package QTLseqR (*77*). All QTLs with an allele frequency difference greater than the 95% CI compared to simulated data which included more than one SNP were retained as candidate loci. This represented a mean cutoff level of 0.143 (i.e. the observed mean allele frequency difference between pools). Note that this cutoff is based on the mean of the smoothed values obtained from QTLseq and it is therefore not directly comparable to the CI-based cutoff applied for the generalized linear model. It should be noted that a caveat with the QTLseq analysis is that the model assumes equal sample sizes of pools (e.g. dead larvae + dead pupae and dead embryos are given equal weight despite the approximately 4-fold sample size difference).

### Demographic inference

To infer the demographic history of the SWE and CAT populations of *L. sinapis* we used the previously described population re-sequencing data, consisting of 10 whole-genome sequenced males for each population. For this analysis, SNPs were filtered to obtain the most reliable variants (Table S12). The resulting SNP data set was thinned using vcftools ver. 0.1.16 (*78*), to ensure that SNPs were at least 10 kb apart. This decreases the impact of physical linkage between sites while ensuring that the whole genome is represented. As a final filtering step, we removed all remaining SNPs inside coding sequence to reduce the impact of selection on the demographic inference. The final SNP set consisted of 59,823 variants. We computed the joint minor allele frequency (MAF) spectrum using easySFS (*79*). The parameters in the demographic model were inferred using GADMA ver. 2 (*80*), which employs a genetic algorithm to optimize parameter values. As an engine in the inference we used Moments (*81*), which fits the observed joint MAF spectrum to simulated data using ordinary differential equations. To transform relative values into estimates of *N_e_* and time in generations since divergence, we assumed a callable sequence length of 7,489,125 bp after filtering, based on π = 0.008 (note that this is lower than the observed levels of genetic diversity due to population expansions). The mutation rate was set to 2.9 * 10^-9^ per base pair and generation – an estimate from a pedigree-based analysis in *Heliconius melpomene* (*82*). Two demographic models were inferred (Isolation-with-migration and Isolation-without-migration) and compared using the Akaike information criterion. Confidence intervals for demographic parameters were estimated based on 100 bootstrap replicates of the joint MAF using the Godambe information criterion (*83*).

### Population genetic analyses

We filtered the population resequencing all-sites variant call format file (including variant and invariant sites) based on depth by marking individuals with < 5 and > 25 reads as missing data using BCFtools *filter* (*84*). Population genetic summary statistics (*F_ST_, D_XY_* and π) were estimated using *pixy* (*85*). We used Hudson’s estimator of *F_ST_*, as recommended by Bhatia *et al.* (*86*). All population genetic summary statistics were estimated in 10 kb genomic windows for three sets of windows: genome-wide (all windows), hybrid incompatibility candidate regions and non-candidate regions. We used ANOVA with a linear model (*X ∼ Chromosome + Type*, where *Type* signifies candidate and non-candidate regions and *X* represents *F_ST_, D_XY_* and π_SWE_ and π_CAT_, respectively) to determine whether the population genetic summary statistics varied with the type of genomic region, while controlling for chromosomal effects such as faster differentiation on the Z chromosome.

### Estimates of the recombination rates

Recombination rate estimates were obtained from pedigree-based linkage maps from the Swedish and the Catalonian populations (for details see refs: (*33*, *36*)). The genetic distance for each marker pair was divided by the physical distance to calculate the expected number of crossover pairs per megabase pair (centiMorgans/Mb).

### Inference of chromosomal rearrangements

To map chromosomal rearrangements to the Asturian *L. sinapis* genome assembly, we performed pair-wise LASTZ ver. 1.04 (*87*) whole-genome alignments to previously published reference assemblies for a SWE and a CAT male, respectively (*36*). Parameters used for both runs of LASTZ were: *M = 254, K = 4,500, L = 3,000, Y = 15,000, C = 2, T = 2,* and *--matchcount = 10,000*. We used previously available data on polarization of fission and fusion events, which were based on synteny analysis based on eight chromosome-level genome assemblies: two each of SWE and CAT *L. sinapis* as well as the outgroup species *L. reali* and *L. juvernica* (*33*, *36*). For example, if a chromosome is fused in *L. juvernica, L. reali* and SWE *L. sinapis* but unfused in CAT *L. sinapis,* then the rearrangement was inferred to be a derived fission in the CAT lineage (Fission CAT). Chromosomes which had a shared breakpoint with outgroups *L. juvernica* and *L. reali* were classified as having unknown polarization (*33*, *36*). Sample size of each rearrangement type was: Fusion SWE = 6, Fission CAT = 5, Unknown = 4.

### Genomic resampling methods

We used a resampling method to evaluate the association between candidate regions for hybrid inviability and other sets of genomic features using a custom script. Chromosomes were randomly chosen, weighted by the length. Coordinates within chromosomes were sampled according to the length of the testing set Y and the overlap between testing set X and reference set Y was calculated. We calculated a two-tailed empirical *p-*value as 2*r n* for *r n* <= 0.5 and 2 (1 -*r/n*) for *r n* > 0.5, where *r* is the number of replicates with an overlap greater than or equal to the overlap for the observed data (*88*). Enrichment was defined as the following odds ratio for two sets; X and Y:

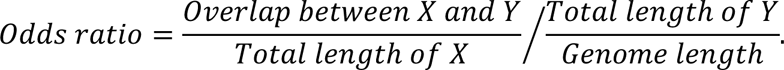

## Data access

DNA-sequencing data is available at the European Nucleotide Archive under study id ERP154226. Scripts will be available in the Github repository: https://github.com/JesperBoman/Evolution-of-hybrid-inviability-associated-with-chromosome-fusions.

## Acknowledgements

This work was funded by the Swedish Research Council (VR research grant #019-04791 to N.B.) and by NBIS/SciLifeLab long-term bioinformatics support (WABI). R.V. was supported by grants PID2019-107078GB-I00 and PID2022-139689NB-I00 funded by MCIN/AEI/ 10.13039/501100011033 and ERDF A way of making Europe, and by grant 2021-SGR-00420 funded by Generalitat de Catalunya. Sequencing was performed by the SNP&SEQ Technology Platform in Uppsala. The facility is part of the National Genomics Infrastructure (NGI) Sweden and Science for Life Laboratory. The SNP&SEQ Platform is also supported by the Swedish Research Council and the Knut and Alice Wallenberg Foundation. We thank Varvara Paida for help developing laboratory methods and Lars Höök, Mahwash Jamy and Arild Husby for valuable input to this project.

