## Supplementary Information for "Evolution of hybrid inviability associated with chromosome fusions"

**Figure S1**: A pedigree tracing the ancestry of the F_2_ population to the F_0_ founders. Circles represent individuals and rectangles represent groups of individuals. For individuals, an identification code is provided, while digits for groups represent the number of females (orange), males (blue) and non-sexed (green) F_2_ offspring for each cross.

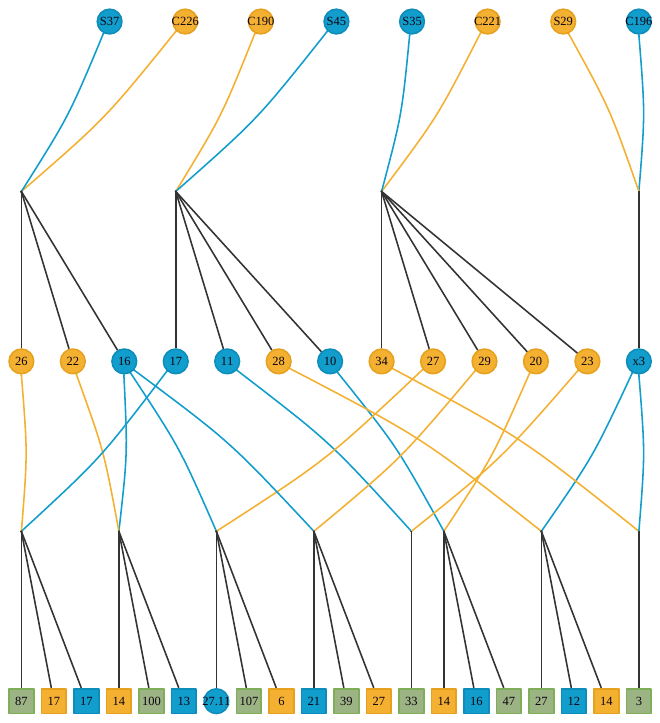

**Table S1**. Survival model. Uninformative prior (*V* = 1, *nu* = 1^-6^). 100,000 iterations. 10,000 iterations discarded as burn-in. Thinning interval = 100, yielding 900 samples.

| **Variable** | **Posterior mean**  **[95 % credible interval]** | **Effective sample size** | **pMCMC** |
| --- | --- | --- | --- |
| Intercept | -0.7568 [-2.2180 – 0.8066] | 900 | 0.322 |
| Animal | 3.08 [1.073 – 5.166] | 526.7 | N/A |

**Table S2**. Survival model. Parameter-expanded prior (*V* = 1, *nu* = 1, *alpha.mu* = 0, *alpha.V* =1000). 100,000 iterations. 10,000 iterations discarded as burn-in. Thinning interval = 100, yielding 900 samples.

| **Variable** | **Posterior mean**  **[95 % credible interval]** | **Effective sample size** | **pMCMC** |
| --- | --- | --- | --- |
| Intercept | -0.7836 [-2.4437 – 0.7604] | 900 | 0.331 |
| Animal | 3.384 [1.497 – 5.756] | 761.5 | N/A |

**Table S3**. Development time model with parameter-expanded priors*. Using Z-scores of development times. 100,000 iterations. 10,000 iterations discarded as burn-in. Thinning interval 100, yielding 900 samples. Family Gaussian.

| **Variable** | **Posterior mean**  **[95 % credible interval]** | **Effective sample size** | **pMCMC** |
| --- | --- | --- | --- |
| Intercept | -1.72100 [-2.19377 – -1.20066] | 343.9 | < 0.001 |
| SexMale | -0.02072 [-0.08679 – 0.03877] | 900.0 | 0.13111 |
| SurvivalAlive | -0.27237 [-0.44827 – -0.11510] | 587.0 | < 0.001 |
| Animal | 0.01061 [8.873e-08– 0.04328] | 97.52 | N/A |
| Units | 0.00114 [0.01459– 0.01771] | 900 | N/A |

* Parameter-expanded prior command*: list(R = list(V = 1, nu = 0.002), G = list(G1 = list(V = diag(6), nu = 0.002, alpha.mu = rep(0, 6), alpha.V= diag(1, 6, 6)), G2 = list(V = 1, nu = 0.002, alpha.mu = 0, alpha.V = 1)))*

**Table S4**. Development time model with uninformative prior*. Using Z-scores of development times. 100,000 iterations. 10,000 iterations discarded as burn-in. Thinning interval 100, yielding 900 samples. Family Gaussian.

| **Variable** | **Posterior mean**  **[95 % credible interval]** | **Effective sample size** | **pMCMC** |
| --- | --- | --- | --- |
| Intercept | -1.31226 [-1.57881– -0.99039] | 16.71 | < 0.001 |
| SexMale | -0.02283 [-0.08696 – 0.04056] | 900 | 0.467 |
| SurvivalAlive | -0.31697 [-0.46309 – -0.18015] | 900 | < 0.001 |
| Animal | 0.04992 [0.0001489- 0.07947] | 13.32 | N/A |
| Units | 0.01616 [0.01466 – 0.01791] | 241.6 | N/A |

* Uninformative prior command*: list(R = list(V = 1, nu = 1e-6), G = list(G1 = list(V = diag(6), nu = 1e-6), G2 = list(V = 1, nu = 1e-6)))*

**Figure S2**: The narrow sense heritability (*h^2^*) distributions for the additive survival model. (A) Using uninformative priors. (B) Using parameter-expanded priors. Both prior specifications showed similar results.

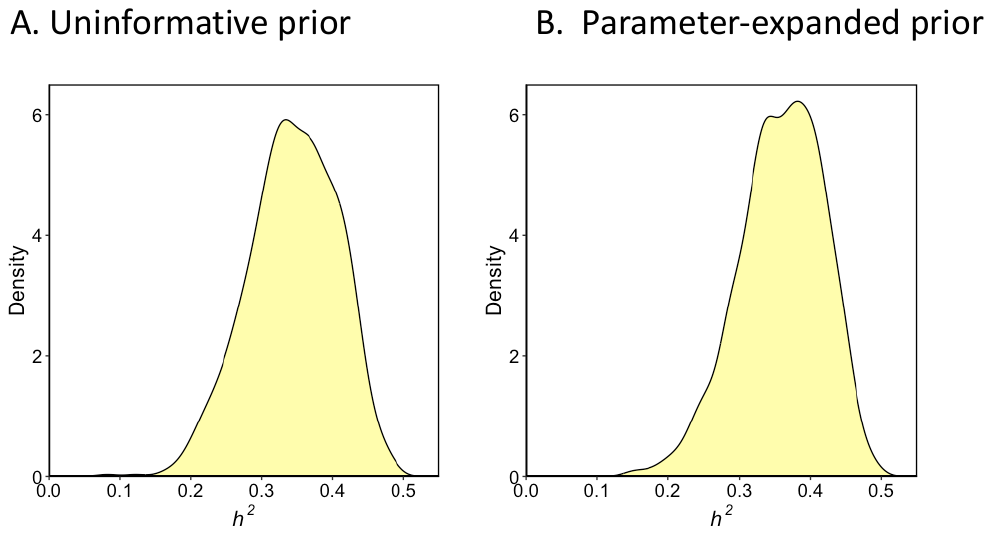

**Table S5**. Summary of sequencing statistics for the pool-seq libraries

| **Pool** | **Sample size** | **Million reads** | **Percentage phred score**  **>=Q30(%)** | **Average**  **read depth with MQ>20*** |
| --- | --- | --- | --- | --- |
| Egg pool | 73 | 393.26 | 91.38 | 115x |
| Dead embryos | 298 | 391.33 | 91.5 | 119x |
| Dead larvae and pupae | 72 | 383.13 | 91.33 | 115x |
| F_2_ adult females | 80 | 398.72 | 91.24 | 115x |
| F_2_ adult males | 76 | 418.98 | 91.61 | 137x |

*At fixed difference marker loci. MQ = mapping quality

**Figure S3.** Allele frequency differences between the *Alive* and *Dead* pools as inferred by QTLseq. Blue lines are the 90 %, 95 % and 99 % CI:s as determined by simulation. Yellow and red boxes represent regions were *Alive* has an excess of SWE and CAT alleles respectively. Excess was defined by having a smoothed allele frequency difference greater than the 95 % CI.

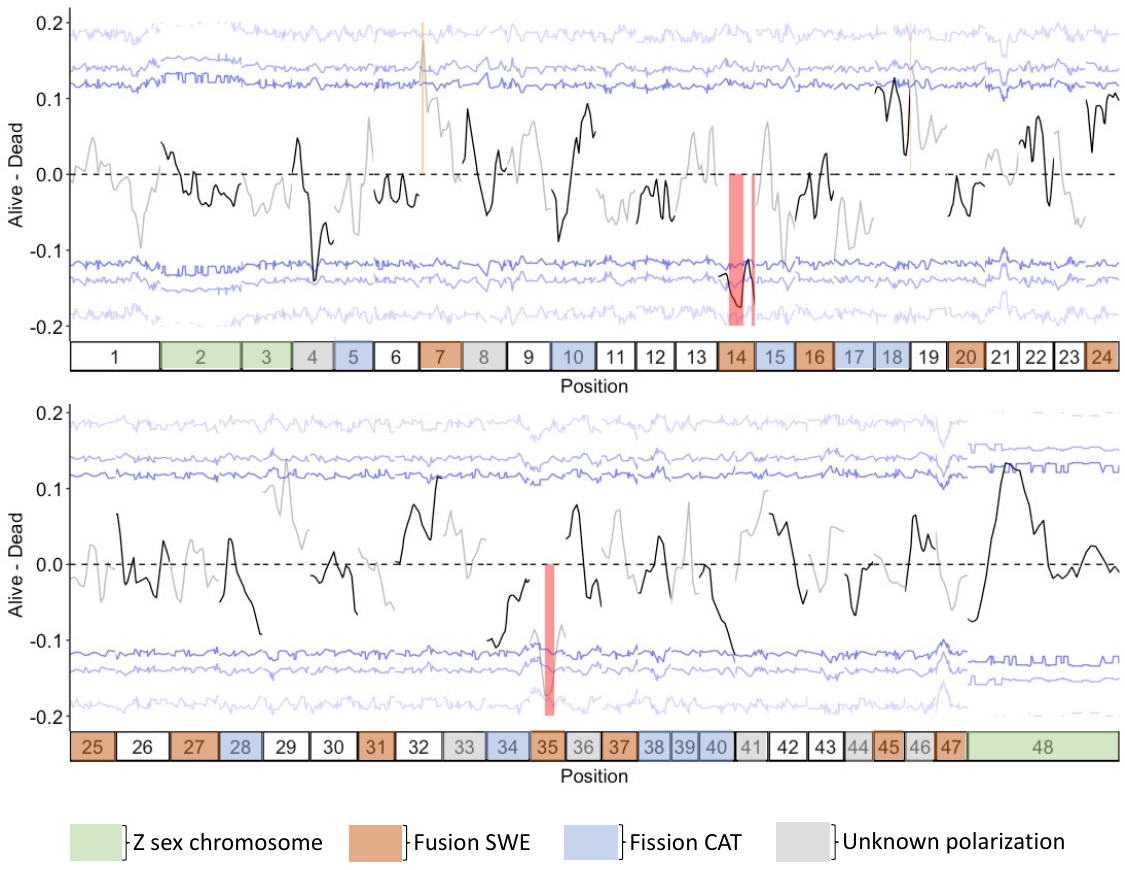

**Figure S4.** Allele frequency differences between the *Alive* pool and the egg pool. The resulting candidate regions significantly overlapped candidate regions obtained from the *Alive* vs *Dead* analysis. Loci have been polarized for the SWE allele frequency. Yellow and red boxes represent regions where the *Alive* pool has an excess of SWE and CAT alleles, respectively. The purple curve shows a generalized additive model that was fitted to the allele frequency differences between genomic regions. We defined candidate regions as stretches of chromosomes where the 95 % CI of the trajectory of the generalized additive model did not overlap an absolute allele frequency difference of 0.075. Chromosomes are plotted on a scale from first to last marker for each individual chromosome. Chromosomes 2, 3 and 48 are the Z-chromosomes. Chromosomes are ordered by size except for chromosome 48 which contains the ancestral Z-chromosome of Lepidoptera. Allele frequencies on Z-chromosomes were normalized by the sample sex ratio. The colors of chromosomes indicate if they represent derived fusions in the SWE population (brown), derived fissions in the CAT population (blue), or segregating fission/fusion polymorphisms (grey). Note that only simple rearrangements (involving two unfused elements) are shown.

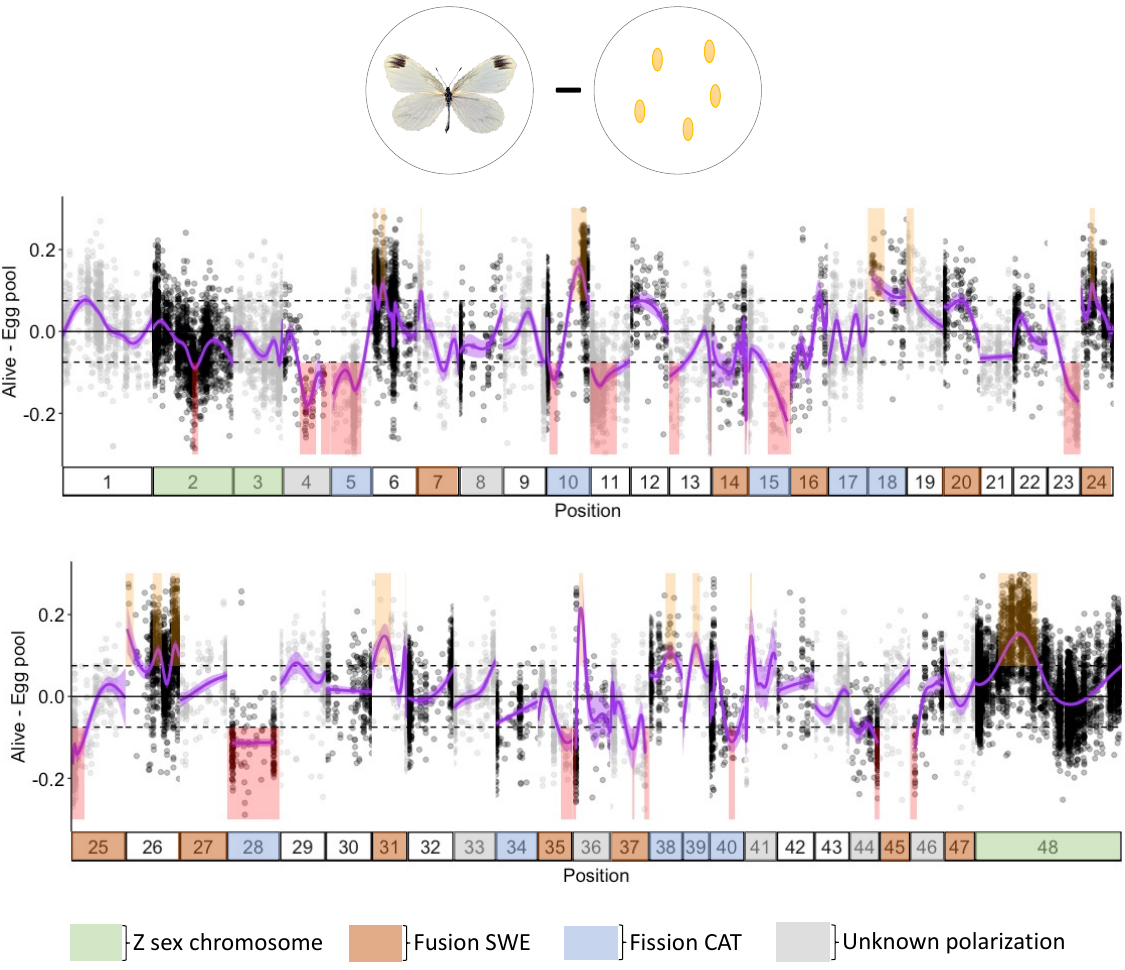

**Table S6.** Associations between chromosomal rearrangements and candidate regions for hybrid inviability when using the QTLseq method to call candidate regions. The analysis was performed for the entire chromosomes, evolutionary breakpoint regions (EBRs) and non-EBR ends of chromosomes, respectively.

| **Type** | **Polarization** | **Odds ratio** | ***p-*value** | ***p-*value*** |
| --- | --- | --- | --- | --- |
| Chromosome | Fission CAT | 0 | <0.001 | <0.001 |
| Chromosome | Fusion SWE | 4.32 | <0.001 | <0.001 |
| Chromosome | Unknown | 0 | <0.001 | <0.001 |
| EBR | Fission CAT | 0 | <0.001 | <0.001 |
| EBR | Fusion SWE | 0 | <0.001 | <0.001 |
| EBR | Unknown | 0 | <0.001 | <0.001 |
| non-EBR ends | Fission CAT | 0 | <0.001 | <0.001 |
| non-EBR ends | Fusion SWE | 0 | <0.001 | <0.001 |
| non-EBR ends | Unknown | 0 | <0.001 | <0.001 |

*Corrected for multiple testing using the Bonferroni method for each category separately.

**Table S7.** Relationship between chromosome fusions and coding sequence density compared to the genome-wide average. Chromosomes involved in derived fusions in the SWE lineage has significantly less coding sequence than the genome on average. This means that the association between hybrid inviability candidate regions and fusions cannot be explained by both being associated with coding sequence density.

| **Type** | **Polarization** | **Odds ratio** | ***p-*value** |
| --- | --- | --- | --- |
| Chromosome | Fusion SWE | 0.932415 | <0.001 |
| EBR | Fusion SWE | 0.658493 | 0.07 |

**Figure S5**: Mean coverage per chromosome in the dead embryo pool (A) and surviving males (B) and females (C). Z sex chromosomes are marked as yellow and autosomes as purple. No chromosomes show indication of systematic aneuploidy (e.g. consistently trisomic) in any pool.

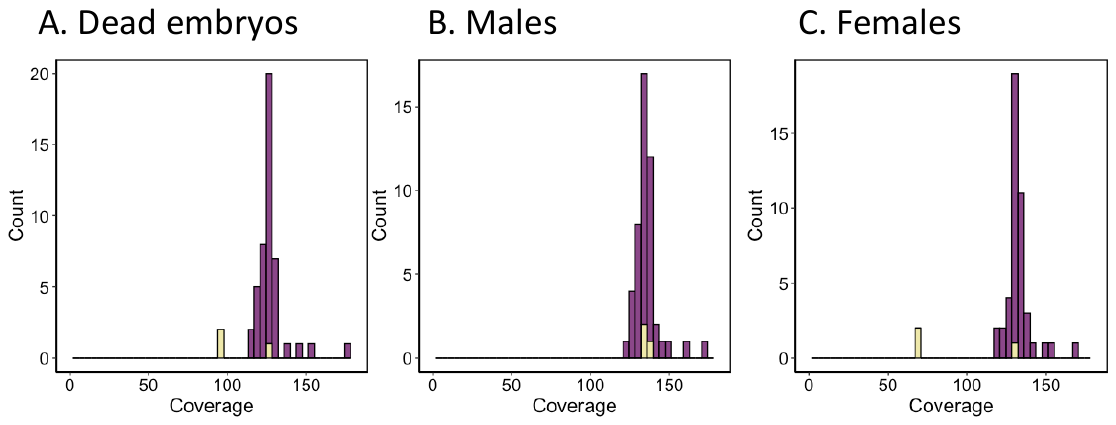

**Table S8.** Average read coverage per chromosome at fixed differences for the dead embryo pool and surviving F_2_ males and females.

| Chromosome | Dead embryos | Males | Females | Chromosome type |
| --- | --- | --- | --- | --- |
| 1 | 126 | 130 | 134 | A |
| 2 | 95 | 68 | 135 | Z |
| 3 | 127 | 130 | 134 | Z |
| 4 | 120 | 127 | 129 | A |
| 5 | 129 | 137 | 143 | A |
| 6 | 139 | 152 | 148 | A |
| 7 | 120 | 126 | 131 | A |
| 8 | 126 | 132 | 133 | A |
| 9 | 124 | 129 | 132 | A |
| 10 | 123 | 129 | 134 | A |
| 11 | 127 | 131 | 136 | A |
| 12 | 129 | 133 | 137 | A |
| 13 | 126 | 138 | 138 | A |
| 14 | 125 | 130 | 131 | A |
| 15 | 126 | 130 | 138 | A |
| 16 | 132 | 140 | 141 | A |
| 17 | 154 | 169 | 173 | A |
| 18 | 113 | 119 | 121 | A |
| 19 | 127 | 132 | 134 | A |
| 20 | 128 | 135 | 136 | A |
| 21 | 175 | 180 | 161 | A |
| 22 | 125 | 134 | 132 | A |
| 23 | 126 | 133 | 137 | A |
| 24 | 124 | 129 | 133 | A |
| 25 | 127 | 132 | 137 | A |
| 26 | 132 | 136 | 139 | A |
| 27 | 129 | 135 | 137 | A |
| 28 | 125 | 135 | 134 | A |
| 29 | 122 | 133 | 135 | A |
| 30 | 123 | 123 | 126 | A |
| 31 | 128 | 129 | 132 | A |
| 32 | 128 | 134 | 138 | A |
| 33 | 128 | 132 | 133 | A |
| 34 | 120 | 125 | 128 | A |
| 35 | 145 | 148 | 147 | A |
| 36 | 126 | 132 | 135 | A |
| 37 | 129 | 129 | 134 | A |
| 38 | 130 | 132 | 131 | A |
| 39 | 123 | 128 | 129 | A |
| 40 | 122 | 129 | 135 | A |
| 41 | 117 | 121 | 125 | A |
| 42 | 125 | 131 | 137 | A |
| 43 | 115 | 133 | 135 | A |
| 44 | 126 | 131 | 133 | A |
| 45 | 120 | 124 | 128 | A |
| 46 | 124 | 129 | 134 | A |
| 47 | 127 | 133 | 132 | A |
| 48 | 97 | 69 | 138 | Z |

**Figure S6**: Parental population recombination rates in candidate regions for hybrid inviability. (A) Recombination rates measured in centiMorgan per Mb (cM/Mb; extracted from pedigree-based linkage-maps) for all candidate regions. Lines connect the recombination rate values of each candidate region. Hair-cross symbols represent the genome-wide recombination rate in each population. Asterisks (*) indicate that recombination rates in the candidate regions for hybrid incompatibility are significantly different from the genome-wide average as determined by resampling.

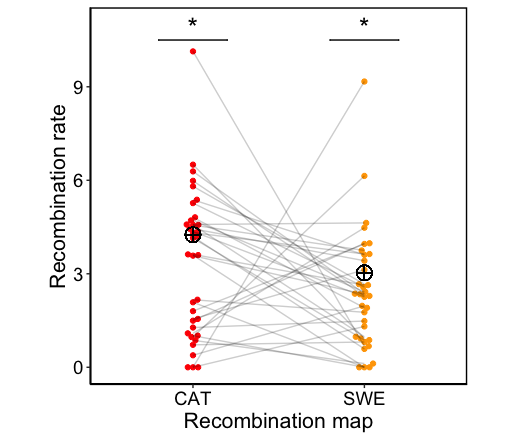

**Table S9.** Recombination rates at fusion EBRs (±1 Mb of breakpoint) compared to genome-wide rates and non-EBR chromosome ends of chromosomes involved in rearrangements, using a SWE and CAT parental population recombination map respectively. EBRs show significantly lower recombination rates than genome-wide but are not significantly different from non-EBRs, regardless of recombination map. Significant values are highlighted in bold.

| **Test** | **Polarization** | **Recombination map** | ***p*** |
| --- | --- | --- | --- |
| EBR vs genome-wide | Fusion SWE | SWE | **0.02009** |
| EBR vs genome-wide | Fusion SWE | CAT | **0.02833** |
| EBR vs non-EBR | Fusion SWE | SWE | 0.2612 |
| EBR vs non-EBR | Fusion SWE | CAT | 0.6699 |

**Figure S7:** Patterns of recombination at fissions and rearranged chromosomes with unknown polarization. Evolutionary breakpoint regions (EBRs) are shown in purple and non-EBR chromosome ends are shown in green. Patterns of average parental recombination rates in EBRs and non-EBRs chromosome ends are presented for 2, 3, 4 and 6 Mb windows. Error bars represent the standard error of the mean. Solid and dashed lines show the recombination rates in the CAT and the SWE population, respectively. Horizontal lines represent mean genome-wide recombination rates for the CAT (red) and SWE (orange) population.

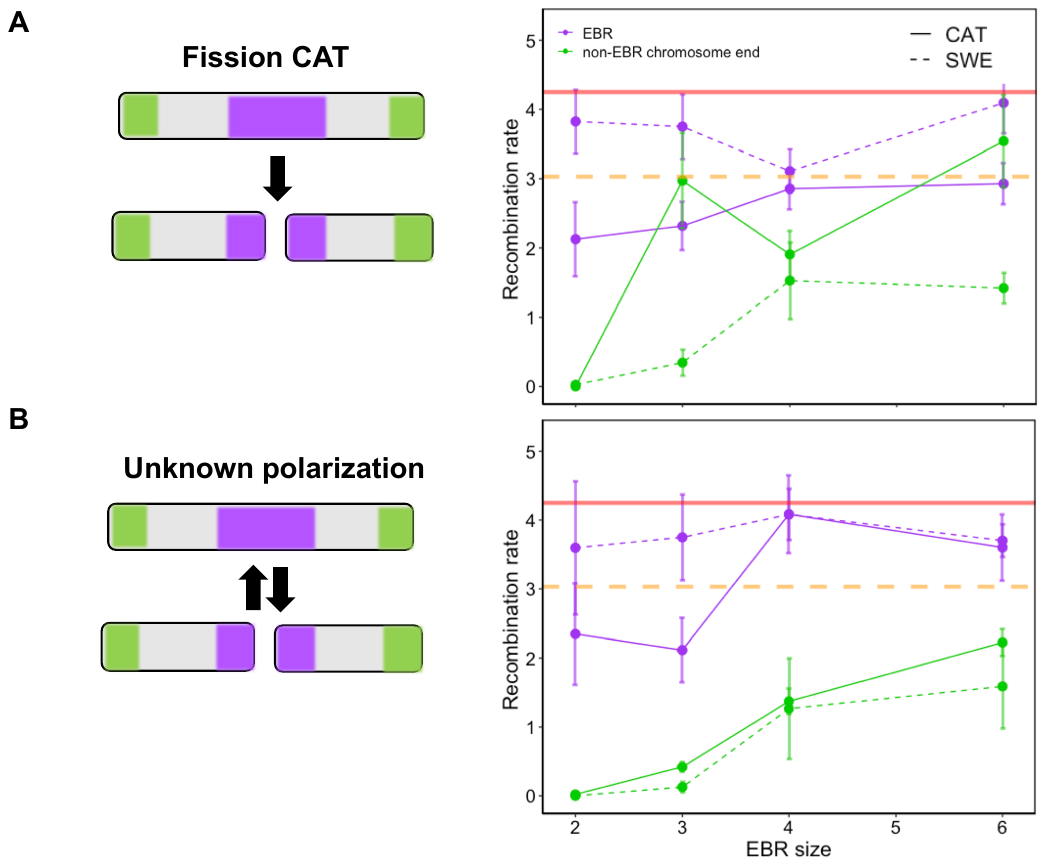

**Table S10.** Recombination rates at EBRs (±1 Mb of breakpoint) of fissions and rearranged chromosomes with unknown polarization, compared to genome-wide rates and non-EBR chromosome ends of chromosomes involved in rearrangements, using a SWE and CAT parental population recombination map respectively. Recombination rates at EBRs for these chromosomes are roughly equal to genome-wide rates but generally higher than their more recombinationally quiescent non-EBR ends. Significant values are highlighted in bold.

| **Test** | **Polarization** | **Recombination map** | ***p*** |
| --- | --- | --- | --- |
| EBR vs genome-wide | Fission CAT | SWE | 0.4922 |
| EBR vs genome-wide | Unknown | SWE | 1 |
| EBR vs genome-wide | Fission CAT | CAT | 0.2685 |
| EBR vs genome-wide | Unknown | CAT | 0.2785 |
| EBR vs non-EBR | Fission CAT | SWE | **0.001163** |
| EBR vs non-EBR | Unknown | SWE | **0.005494** |
| EBR vs non-EBR | Fission CAT | CAT | 0.06639 |
| EBR vs non-EBR | Unknown | CAT | 0.1956 |

**Table S11**: Inference of the demographic history of SWE and CAT *L. sinapis.* Two models were fit: an isolation model assuming no migration since divergence and isolation-with-migration. The latter model provided a superior fit as evaluated by AIC. Numbers in brackets show the 95 % confidence interval for relevant parameters.

|  | **Isolation** | **Isolation-with-migration** |
| --- | --- | --- |
| **AIC** | 6218 | 2480 |
| **Number of parameters** | 4 | 6 |
| **Log likelihood** | -3105.29 | -1245.94 |
| ***T*** | 0.18 [-0.1 – 0.3] | 403,329 [112,035 – 672,215] |
| **Proportion of split** | 0.86 [0.8 –1.0] | 0.13 [-0.1 – 0.4] |
| **N_ANC_** | 354,588 | 280,418 |
| **N_CAT_** | 450,792 [35,589 – 957,387] | 578,270 [196,500 – 954,426] |
| **N_SWE_** | 1,924,429  [1,560,187 – 2,269,363] | 396,180 [112,391 – 674,349] |
| **M_SWE🡪CAT_** | NA | 1.07 [0.5 –1.6] |
| **M_CAT🡪SWE_** | NA | 0.18 [0 – 0.4] |

**Figure S8.** Relationship between absolute divergence (*D_XY_*) and genetic differentiation (*F_ST_*) between SWE and CAT *L. sinapis*. The two measures of genetic difference showed a weak but significant correlation (Spearman’s ρ = 0.11, *p <* 2.2 *10^-16^), indicating that barriers to gene flow has had a mild influence on the genomic landscape of differentiation. Alternatively, the divergence may be too recent for such an effect to be prominently seen (*49*).

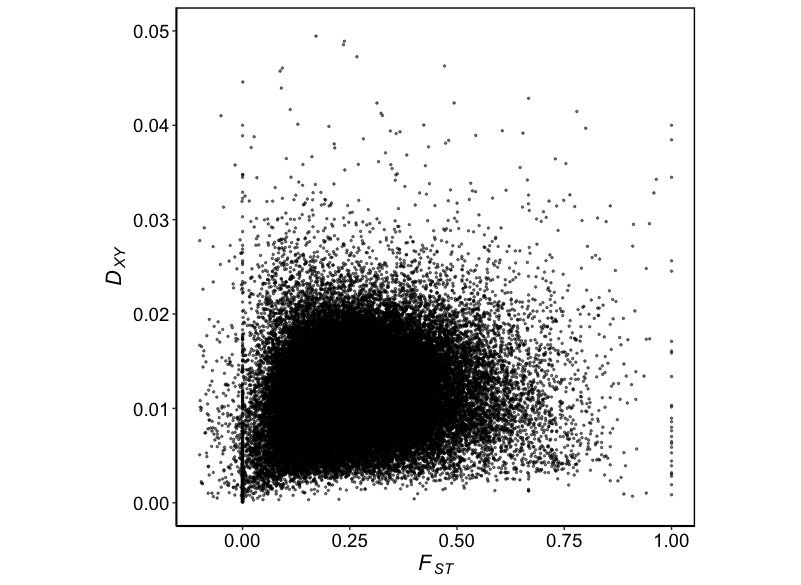

**Figure S9.** Population genetic summary statistics per chromosome with special emphasis on differences between autosomes and Z sex chromosomes. Z chromosomes have lower average pairwise difference (π), and absolute genetic divergence (*D_XY_*) but higher genetic differentiation (*F_ST_*) compared to autosomes. Bars in boxes represent the median value per chromosome in non-overlapping 10 kb windows. Box hinges are the 25th and 75th percentiles, whiskers extend to 1.5 times the distance between the 25th and 75th percentiles and solid dots represent outliers.

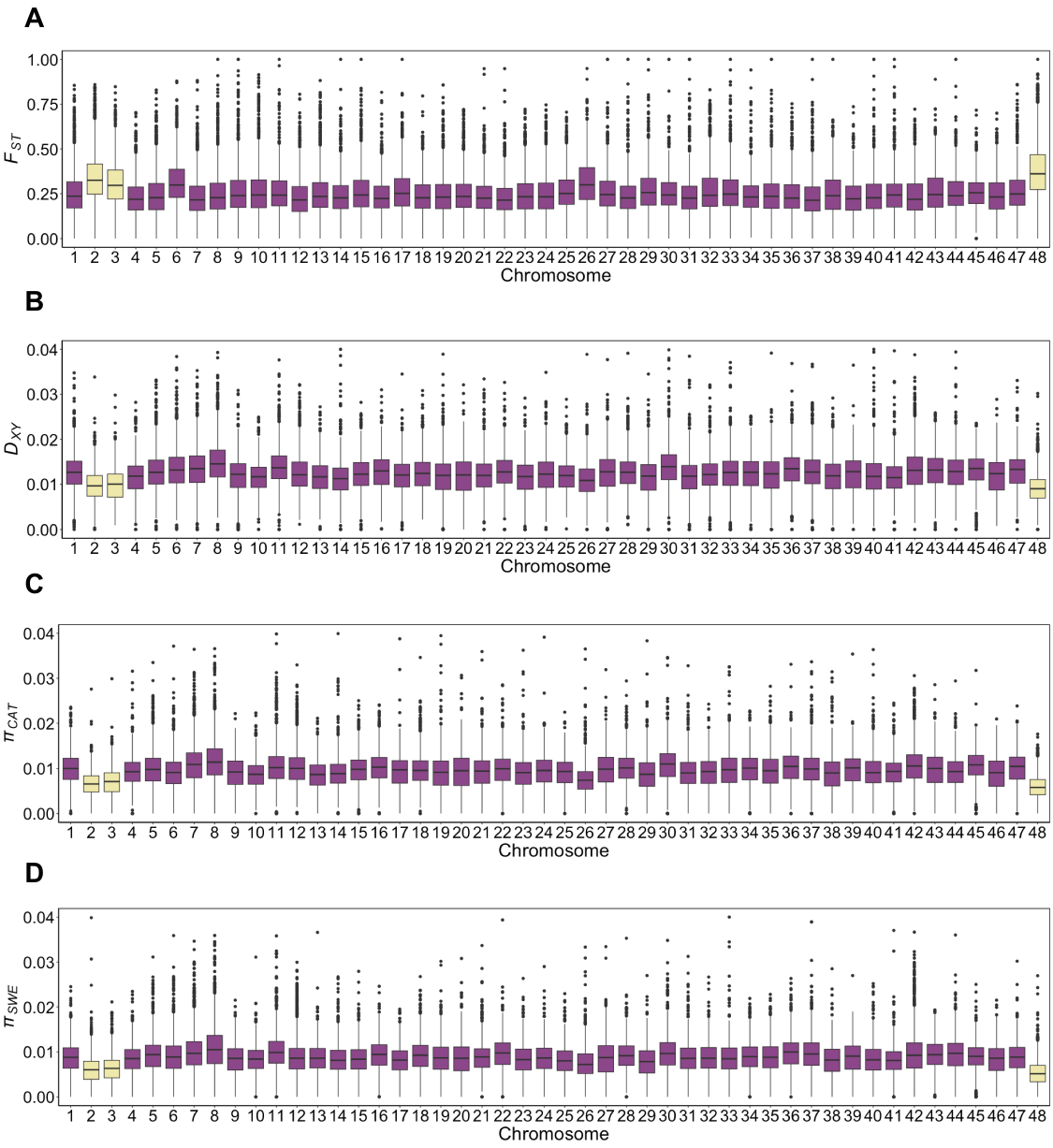

**Figure S10:** Population genetic summary statistics across all chromosomes measured in non-overlapping 10 kb windows. Values have been Z-transformed (subtracted by the mean and divided by the standard deviation) for ease of illustration. In addition, local regression curves (span=0.01, degree=1) were fit to the data, with shaded regions representing 95 % confidence intervals. Genetic differentiation (*F_ST_*) is shown in blue, absolute divergence (*D_XY_*) in dark green*,* genetic diversity in the CAT population (π_CAT_) in red and SWE (π_SWE_) in orange respectively. Boxes represent candidate regions for hybrid incompatibility between SWE and CAT *L. sinapis*. Of note, large peaks of *D_XY_* and π on chromosomes 11, 12 and 42 possibly represent collapsed assembly regions due to an increased coverage among some *L. sinapis, L. juvernica* and *L. reali* (data not shown).

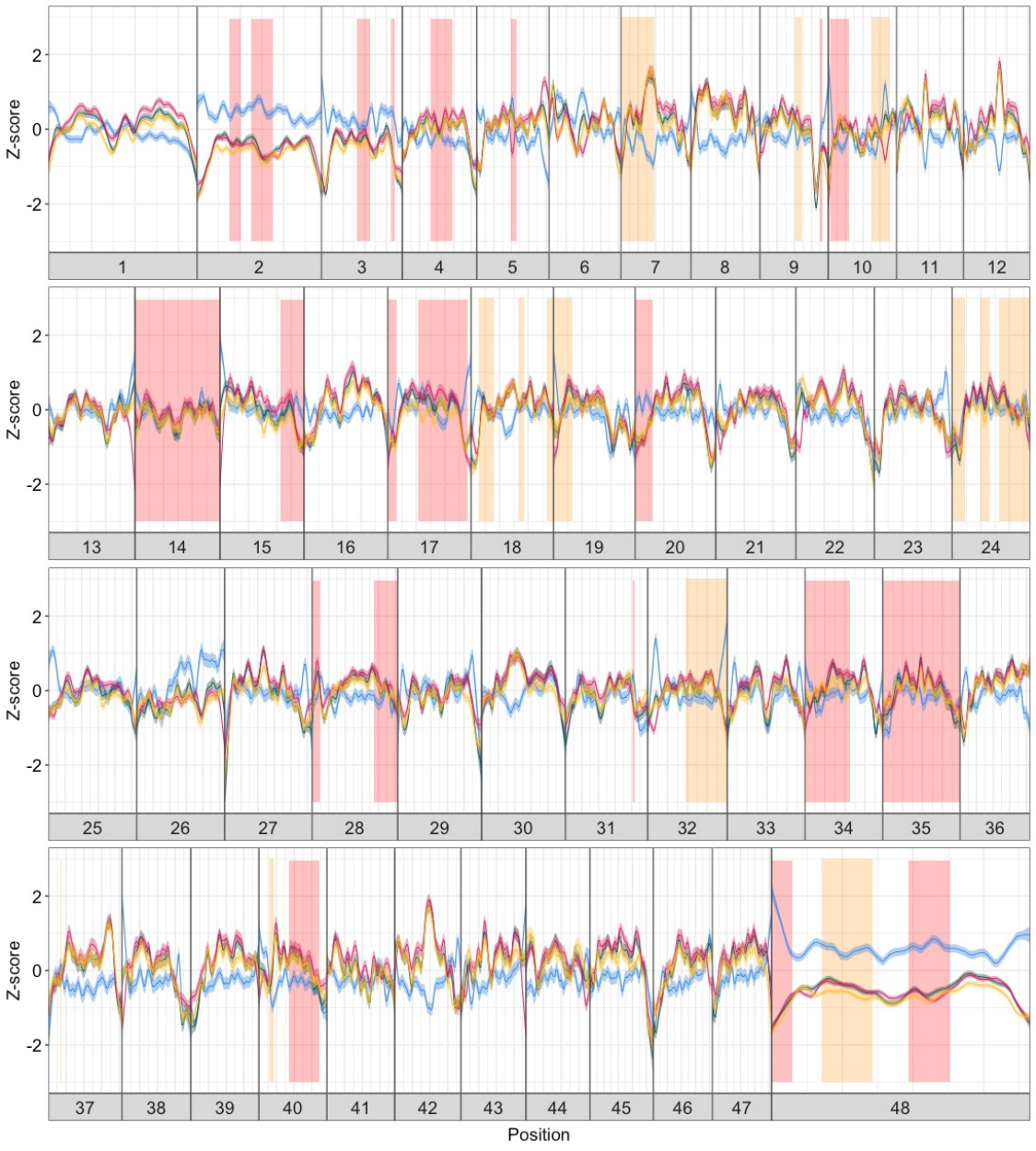

**Table S12**. Filtering parameters for population-resequencing data to obtain a set of high-quality SNPs.

| **Parameter** | **Filtering threshold** |
| --- | --- |
| Fisher strand bias | <60 |
| Strand odds ratio | <3 |
| Mapping quality | >40 |
| Mapping quality rank sum test | >-12.5 |
| Quality by depth | >2 |
| Read position rank sum test | >-8 |
| Depth | <31.09 (3 standard deviations above mean read coverage) |
| Max missing | 1 (no missing allele information) |
